# Early posttranscriptional response to tetracycline exposure in a gram-negative soil bacterium reveals unexpected attenuation mechanism of a DUF1127 gene

**DOI:** 10.1101/2025.01.31.635925

**Authors:** Jennifer A.F. Kothe, Till Sauerwein, Theresa Dietz, Robina Scheuer, Muhammad Elhossary, Susanne Barth-Weber, Jan Wähling, Konrad U. Förstner, Elena Evguenieva-Hackenberg

**Affiliations:** Institute of Microbiology and Molecular Biology, University of Giessen, 35392 Giessen, Germany; ZB MED - Information Center for Life Sciences and University of Cologne, 50931 Cologne, Germany; TH Köln – University of Applied Sciences, 50678 Cologne, Germany

## Abstract

The Gram-negative, soil-dwelling plant symbiont *Sinorhizobium meliloti* shares its free-living habitat with antibiotic producers. To learn about early steps of its adaptation to antibiotics, we analyzed transcriptome changes after 10 min exposure to subinhibitory amount of tetracycline (Tc). RNA-seq revealed 297 differentially expressed genes. Besides ten upregulated ribosomal genes, there was no recognizable functional pattern in the observed changes. Instead, upregulated genes were mostly first in operons and often small, while downregulated genes were downstream in operons. Furthermore, we detected mRNA stabilization upon Tc exposure for several up- and down-regulated genes. Thus, mRNA stabilization contributed to increased mRNA levels, but for downstream genes its effect was counteracted by premature transcriptional termination caused by disrupted coupling between transcription and translation. Using reporter constructs, we found that a DUF1127 gene, showing highest mRNA increase, is controlled by transcription attenuation depending on the translation of an upstream ORF (uORF). Our data suggest the following model: The attenuation strongly depends on the accessibility of C-rich codons at the begin of the uORF. The accessibility is guaranteed by translation of the uORF, and is possible in a time window after a ribosome moves downstream and before a next ribosome occupies the ribosomal binding site (RBS). The accessibility is blocked either by impaired translation initiation or, in the absence of ribosome binding, by base-pairing between the RBS and the C-rich codons. We propose that this is used by bacteria to monitor ribosome availability and translation efficiency, and to ensure reciprocal expression of the DUF1127 gene.

## Introduction

Soil bacteria share their habitat with many antibiotic producers. Therefore, and also due to (mis)use of antibiotics in agriculture, they are often exposed to antibiotics (Nelson und Levy 2011); (Wang et al. 2024). Adaptation of soil bacteria to antibiotic exposure is important for their survival and competitiveness. Knowledge about the adaptation mechanisms could help to better understand the resistance and tolerance strategies of pathogens. However, for pathogenic and for environmentally important model bacteria, information about very early steps in response to antibiotics is scarce, although it is conceivable that these steps could be decisive for adaptation.

Upon antibiotic exposure, transcription and/or translation of resistance genes (for example encoding efflux pumps or rRNA modifying enzymes) is induced (Hahn et al. 1982); (Dersch et al. 2017); (Weston et al. 2018); (Kesavan et al. 2020); (Urban-Chmiel et al. 2022); (Takada et al. 2022). However, even sub-inhibitory antibiotic amounts lead to phenotypic changes (Morita et al. 2014) and transcriptome studies revealed that many genes are affected in the treated bacteria. For example, the mRNA levels of hundreds of genes were differentially regulated in a pathogenic *Pseudomonas aeruginosa* after 2 h of exposure to inhibitory amounts of tobramycin or colistin. Specifically, the translation-inhibiting antibiotic tobramycin led to differential regulation of genes involved in translation, amino acid catabolism, central metabolism, and secretion (Cianciulli Sesso et al. 2021). On the other hand, 10 min exposure of *Acinetobacter baumannii* to subinhibitory amount of the translation-inhibiting antibiotic minocycline resulted in changes in mRNA levels of only 25 protein-coding genes, among them stress response and transport genes (Gao und Ma 2022). While in these studies the molecular mechanisms of changes in mRNA levels were not studied, it is known that in gram-negative bacteria, inhibition of translation leads to Rho-dependent premature transcriptional termination (PTT). The PTT results from decoupling between translation and transcription (Kohler et al. 2017), and has a negative polar effect on gene expression with increasing distance from the transcriptional start site (Zhu et al. 2019). In *E. coli*, PPT in response to sublethal amounts of the translation-inhibiting antibiotics chloramphenicol and erythromycin was detected for long genes such as *lacZ* and for large operons (Zhu et al. 2019).

Regulated PTT in the mRNA leader, also known as transcription attenuation, is often used by Gram-positive bacteria to keep low the expression of resistance genes in the absence of antibiotics (Dar et al. 2016). This mechanism relies on small upstream open reading frames (uORFs). Typically, efficient translation of an uORF ensures the formation of an intrinsic transcriptional terminator in the nascent RNA. Upon antibiotic exposure, the ribosome stalls at the uORF, the nascent RNA adopts an alternative RNA structure (antiterminator) and the downstream resistance gene is induced (Lee et al. 2022); (Takada et al. 2022). In Gram-negative bacteria, similar ribosome-dependent transcription attenuation is used for regulation of amino acid biosynthesis genes (Keller und Calvo 1979); (Merino et al. 2008). The transcription attenuation can also be Rho-dependent (Turnbough 2019). Rho is a hexameric ATPase that binds a Rho utilization site (*rut*-site; approximately 90 nt region with C>G content) for its action (Hao et al. 2021). Ribosome stalling in an uORF can lead to accessibility of a *rut*-site or of a Rho-antagonizing RNA element (RARE) in the downstream nascent RNA (Ben-Zvi et al. 2019); (Sevostyanova und Groisman 2015).

High-throughput detection of PPT in bacterial 5’-UTRs was achieved by global mapping of 3’-ends and in many cases, uORFs were linked to conditional 3’-ends (Dar et al. 2016), (Adams et al. 2021). Bacterial uORFs (and small protein-encoding genes in general) are still insufficiently annotated in bacterial genomes, particularly due to technical difficulties with the ribosome profiling technology for bacteria (Vazquez-Laslop et al. 2022). We are working with *Sinorhizobium meliloti*, a gram-negative soil-dwelling plant symbiont. In the past, the transcriptome of *S. meliloti* 2011 was intensely studied by RNA-seq (Sallet et al. 2013; Roux et al. 2014). Additionally, a recent ribosome profiling study of this strain revealed new translated small ORFs (Hadjeras et al. 2023). Thus, *S. meliloti* 2011 is well-suited to study how short antibiotic exposure shapes a bacterial transcriptome.

Here, we used RNA-seq to analyze transcriptome changes in *S. meliloti* 2011 cultures that were exposed for ten minutes to subinhibitory amount of tetracycline (Tc), an antibiotic commonly detected in soils (McManus et al. 2002); (Robles-Jimenez et al. 2021), (Chang et al. 2023)). The observed transcriptome changes correlated with the position of genes in operons, suggesting that besides the mRNA stabilization, which we detected for several genes, PTT was a major factor shaping the transcriptome. Furthermore, we found that the gene with highest mRNA increase, one of several upregulated DUF1127 homologs, is regulated by transcription attenuation relying on the translation of an uORF. Our data suggest a double control of the accessibility of a possible *rut* site at the begin of the uORF. The *rut* site contains C-rich codons and is occluded either by a ribosome with impaired translation initiation or, in the absence of ribosome binding, by an RNA secondary structure. This suggests that the DUF1127 gene is upregulated in response to translation inefficiency, which is sensed by its uORF-containing mRNA leader.

## Results

### Transcriptome changes upon Tc exposure reveal polar effects in operons

To investigate early effects of tetracycline (Tc) on the *S. meliloti* 2011 transcriptome, exponentially growing cultures were treated with subinhibitory amount of Tc (1.5 µg/ml) for 10 min. Total RNA of the treated cultures was compared to the RNA of non-treated controls. The analysis revealed **297** differentially expressed genes (DEGs) with log_2_(FC) > 1 and p_adj_ ≤ 0.01. Thereby, the mRNA levels of **127** genes were increased, while for **170** genes a decrease was detected (Fig. 1A; Table S1-a).

**Figure 1.**
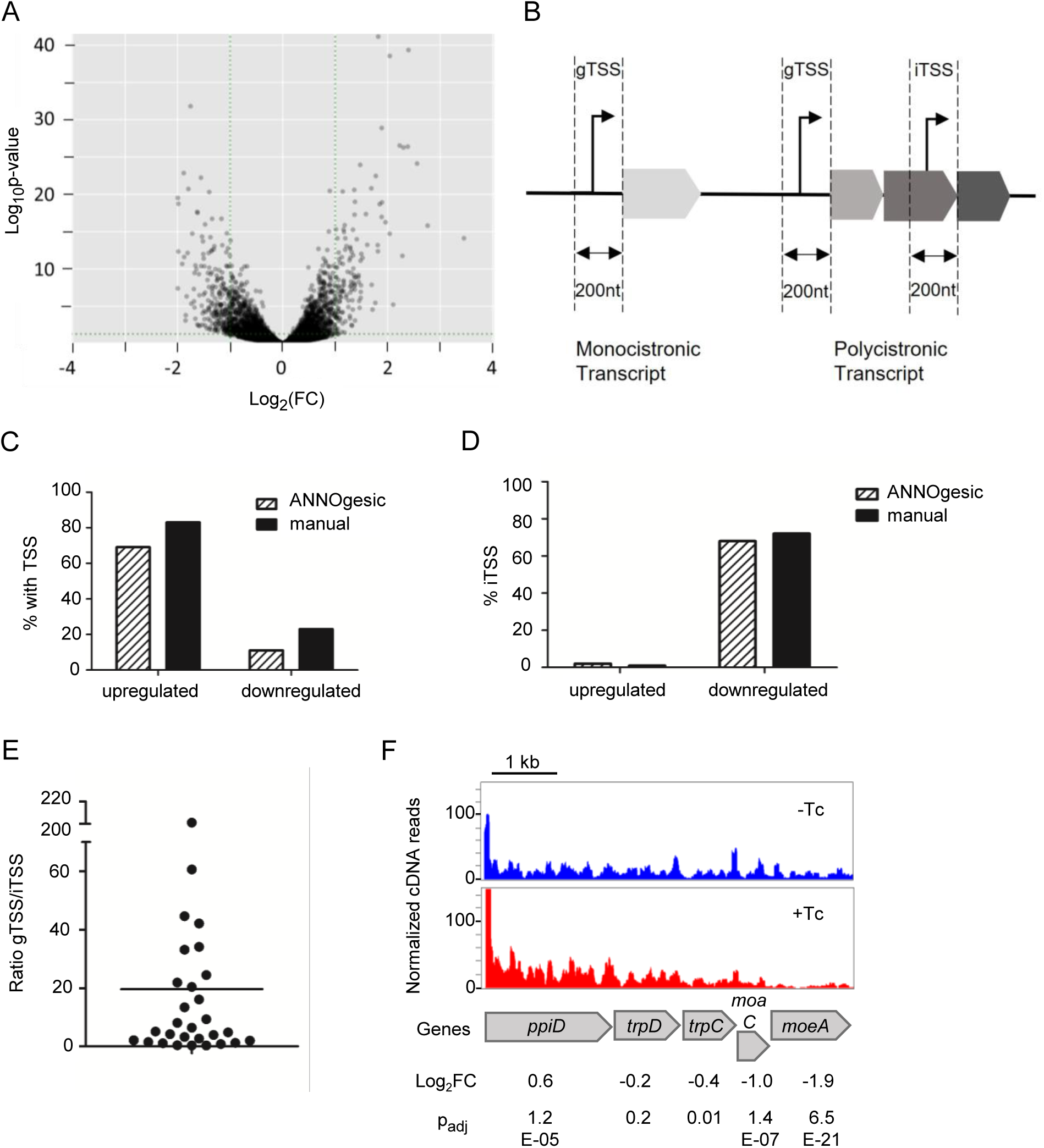
RNA-seq revealed polar effects in *Sinorhizobium meliloti* 2011 operons upon short-term exposure to subinhibitory tetracycline. A) Volcano plot of RNA-seq data showing comparison of Tc-treated cultures (1.5 µg/ml tetracycline for 10 min) to non-treated controls. **B)** Scheme of the transcription start site (TSS) classification used in panels C, D and E. gTSS: gene TSS, a TSS that is not located in an upstream gene and is thus probably not located in an operon. iTSS: internal TSS, a TSS located in an upstream gene and therefore probably located in an operon. Genes with assigned TSSs are indicated in Table S1-b and Table S1-c. **C)** Proportion of upregulated or downregulated differentially expressed genes (DEGs) with TSSs located up to 200 nt upstream of their start codons (gTSSs and iTSSs). TSSs were detected using ANNOgesic (Förstner et al., 2014) and manually. **D)** Proportion of genes with iTSSs among the DEGs with assigned TSSs. **E)** Ratio of the normalized cDNA reads of the gTSS with a highest peak to the corresponding downstream iTSS of a DEG. **F)** Integrated genome browser view of an operon showing RNA increase of the first gene and decrease towards the 3’-end of the polycistronic transcript after Tc addition. Shown are normalized cDNA reads of one of the three biological experiments. The results of the three experiments are indicated below the panel (see Table S1-a) . The first gene (*ppiD*) is long and shows an RNA level increase below the threshold of 2-fold change, but the very low p_adj_ value shows that the increase is significant. The second gene shows no significant change, while the RNA of the downstream genes is decreased. The decrease in mRNA amount is stronger towards the 3’-end of the operon.

We attempted to classify the DEGs based on Gene Ontology (GO). Out of the 127 DEGs with increased mRNA levels (upregulated DEGs), only 23 were in the obtained GO list, and ten of them were assigned to the only significantly enriched GO term “structural constituent of ribosome”. Of the 19 listed GO terms, 16 contain only individual genes (Table S2). Similar results were obtained by a Kyoto Encyclopedia of Genes and Genomes (KEGG) enrichment analysis: Only 35 upregulated DEGs were assigned to KEGG pathways. Highest enrichment (seven genes) was obtained for “ribosome – *Sinorhizobium meliloti*”. Of the 46 listed KEGG pathways, 33 contain only individual genes (Table S3). None of the 170 downregulated DEGs with log_2_(FC) > 1 and p_adj_ ≤ 0.01were assigned to GO terms or KEGG pathways. Besides the ribosome-related genes, the results argue against upregulation or downregulation of specific operons and regulons, and suggest that general and possibly posttranscriptional mechanisms are operating immediately upon Tc exposure.

Visual inspection of cDNA reads using the integrated genome browser (IGB) suggested different localization of upregulated and downregulated DEGs in operons. To address this in more systematic way, we analyzed whether the DEGs are preceded by a transcriptional start site (TSS). To this end, first we used ANNOgesic (Yu et al. 2018) and RNA-seq data for genome-wide TSS mapping in *S. meliloti* cultivated semiaerobically in rich TY medium (the growth conditions used in this work). The mapped TSSs are listed in Table S4. Then we determined for each DEG whether there is a TSS in its -200 region (+1 is the first nucleotide of the start codon). We categorized the TSSs in two groups: The first group comprises TSSs located in upstream genes that are transcribed in the same direction. Such TSSs are most probably internal TSSs (iTSSs) in operons (Fig. 1B). The second group is composed of TSSs that are not located in upstream genes, indicating that they are at the begin of operons or monocistronic transcripts. We designated them regular gene TSSs (gTSSs; (Čuklina et al. 2016)). The presence or absence of a TSS upstream of a DEG is shown in Table S1-b and Table S1-c. We found that most upregulated DEGs are preceded by TSSs, the vast majority of which belongs to the gTSS class (only two iTSS were present; Fig.1C and 1D). By contrast, a minority of downregulated DEGs was found to be preceded by TSSs, and most of them were iTSSs (Fig.1C and 1D). We repeated the analysis manually (see material and methods). While with the manual analysis more TSSs were proposed, the results remained similar: 1) mostly the upregulated DEGs and not the downregulated ones were was preceded by a TSS; 2) The TSSs of the upregulated DEGs were almost exclusively gTSSs; 3) The TSSs of the downregulated DEGs were mostly iTSSs (Table S1-b and Table S1-c; Fig.1C and 1D). Many of the manually proposed iTSSs of downregulated DEGs were weak in comparison to the major gTSS of the respective operon: For 14 out totally 29 iTSSs,the gTSS/iTSS peak ratio was >5 (Fig. 1E).

Thus, analyzing how short Tc exposure shapes the transcriptome of *S. meliloti*, we made the folllowing observations: 1) the majority of the DEGs were not enriched in GO terms or KEGG pathways; 2) upregulated genes are mostly at the begin of operons or in monocistronic transcripts; 3) downregulated genes are mostly second or further downstream in operons. Fig. 1F shows an example of these polar effects in response to short Tc exposure in the *S. meliloti trpDC* operon. The increase in the mRNA level of first genes in transcripts could be explained by mRNA stabilization, while the mRNA decrease of downstream genes in polycistronic transcripts probably reflects PTT caused by decoupling between translation and transcription (Zhu et al. 2019) in the presence of Tc.

### Small genes and DUF1127 genes are overrepresented among the DEGs with highest increase

It is known that in gram-negative bacteria, translation inhibition leads to PPT in long genes (Zhu et al. 2019). Therefore, we predicted that small genes at the begin of operons or in monocistronic transcripts should be overrepresented among the *S. meliloti* DEGs with highest mRNA increase. Indeed, among the 30 genes with highest increase, 13 (43 %) encode proteins shorter than 100 aa (Table S1-b). Furthermore, two of the top 30 upregulated DEGs are preceded by recently identified sORFs (Hadjeras et al. 2023): The sORF3 and sORF55 are upstream of SM2011_RS23660, the first large gene in an ABC-type multidrug transport system operon, and sORF26 is upstream of the gene with highest upregulation upon Tc exposure, which encodes a DUF1127 protein (Table S1). We note that six DUF1127 genes (here designated *duf1127_1_* to *duf1127_6_*) are among the top 30 upregulated DEGs (Table S1-b).

For the gene with the highest upregulation upon Tc exposure, *duf1127_1_* (Table S1-a), we did not detect a TSS with the approach described above (Table S1-b), because the TSS is located 279 nt upstream of its start codon and 30 nt upstream of the 29-aa sORF26 (here referred to as “upstream ORF” or “uORF”; Fig. 2A and Fig. S1). Using Northern blot hybridization, we validated the strong induction of *duf1127_1_* after Tc addition to bacterial cultures and showed the existence of an induced sRNA that contains the uORF (Fig. 2B). Additionally, we validated the RNA-seq data by qRT-PCR analysis of *duf1127_1_* and four other upregulated genes (Fig. 2C). We also tested whether the Tc concentration and the time of treatment played a role in the increase of mRNA levels. For this, *S. meliloti* 2011 was either treated with 0.5, 1 or 1.5 µg/ml Tc for 10 min. A different set of *S. meliloti* cultures was treated with 1.5 µg/ml Tc for 3, 6, or 10 min. RNA isolated from the cultures was analyzed by qRT-PCR, which could verify a dose- and time-dependent response (Fig. 2D and Fig. S2). However, only for *duf1127_1_* statistically significant increase was measured already after 3 min (Fig. 2D).

**Figure 2.**
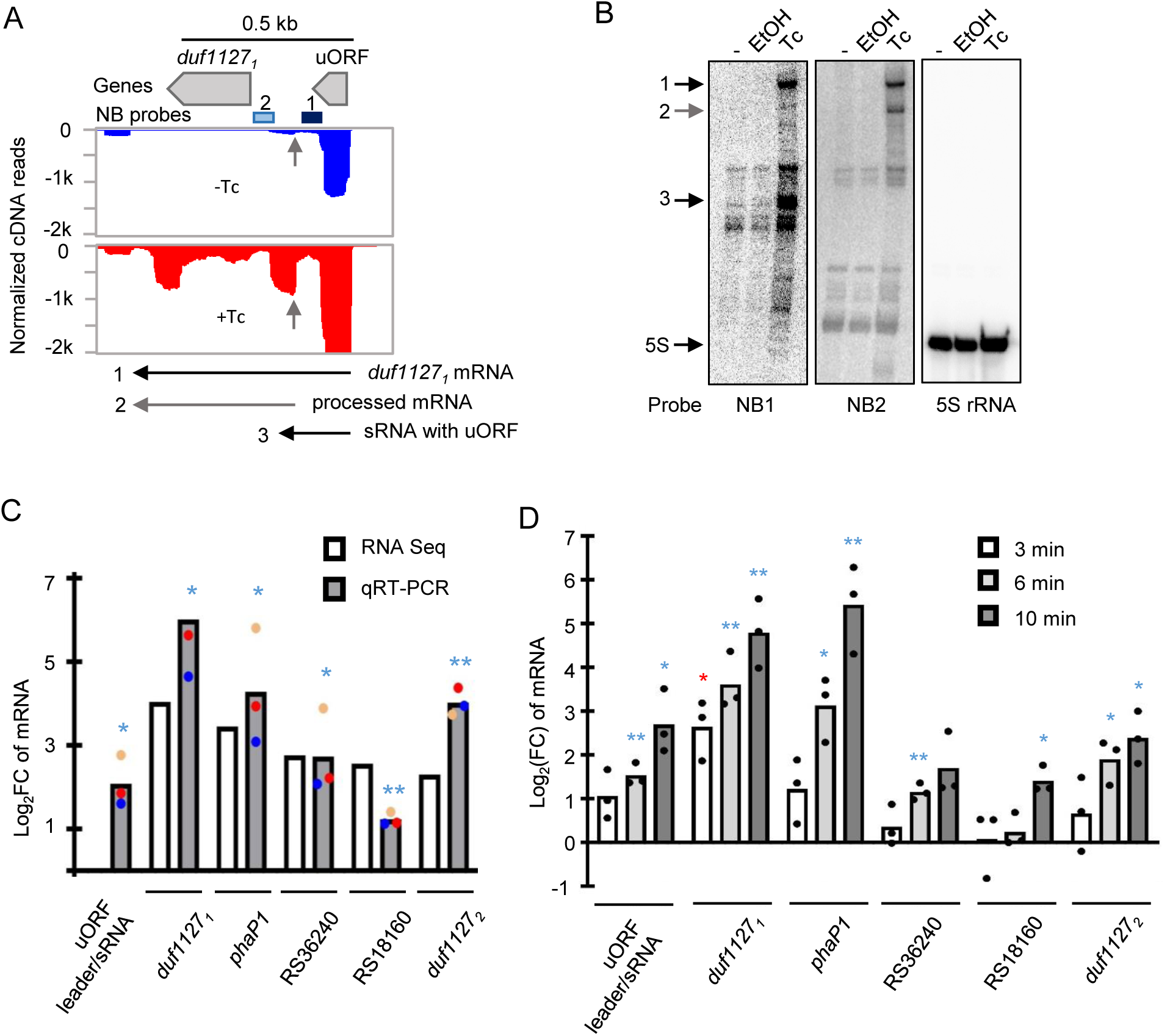
The *duf1127_1_* gene is preceded by an uORF and is strongly upregulated upon Tc exposure. **A)** Integrated genome browser view showing the normalized cDNA reads of one of the three biological experiments. Above the panel, the uORF and the small *duf1127_1_* gene are depicted, along with the position of the oligonucleotides used as probes for Northern blot (NB) hybridization. Below the panel, the main detected transcripts are depicted with horizontal, numbered arrows. In the panel, short vertical arrows mark the 5’-end of a processed transcript. This processing seems not to depend on the tetracycline (Tc) exposure (Fig. S1). **B)** Ten minutes after Tc addition to the cultures, transcripts of the *duf1127_1_* operon were detected by Northern blot hybridization of total RNA separated in 10% polyacrylamide-urea gel. RNA was isolated from three parallel 30 ml-cultures grown to OD_600 nm_ of 0.5. -: no addition of a compound. EtOH: addition of 4.5 µl ethanol. Tc: addition of 4.5 µl ethanol with dissolved Tc (final concentration of 1.5 µg/ml). The same membrane was re-probed and the probes are indicated below the panels. On the left side, the detected induced transcripts are marked with numbered arrows (compare to panel A). 5S rRNA was used as loading control. Shown is a result of a representative experiment. **C)** Validation of the RNA-seq results by qRT-PCR analysis of the indicated genes. The uORF was not annotated and thus not recorded in the table of differentially expressed genes determined by RNA-seq (Table S1-a). **D)** Analysis by qRT-PCR of RNA changes in time after Tc addition. All graphs show means and single data points of three independent experiments. In C), the points of the individual experiments are marked with a specific color. *** p ≤ 0.001, ** p ≤ 0.01, * p ≤ 0.05.

It is noteworthy that for the *duf1127_1_* operon, the Cq of the uORF-containing leader was lower than the Cq of the *duf1127_1_* (ΔCq of approximately 5.5). Upon Tc exposure, the Cq difference became smaller, because the increase of the *duf1127_1_* part of the transcript was stronger than that of the leader (see Fig. 2C, 2D and Fig. S2). This could be explained by different half-life changes of the transcript segments in response to Tc, and/or by transcription attenuation between the uORF and *duf1127_1_*, which is relieved upon Tc exposure.

### mRNA is stabilized upon Tc exposure

Next, we tested whether mRNA stability is changed upon Tc exposure. After stop of transcription by adding rifampicin to *S. meliloti* cultures, mRNA decay was measured using qRT-PCR. In this analysis, we included the highly upregulated genes *duf1127_1_* (and its uORF-containing mRNA leader), *phaP1, duf1127_2_* (Table S1-b), and the first and last genes of the *trpDC* operon (*ppiD* and *moeA*; Fig. 1B and Table S1-a). Additionally, *metZ* was analyzed as an mRNA with a short half-life (Scheuer et al. 2022), the level of which was not changed upon Tc exposure (Table S1-a). In the absence of Tc, the half-life of *metZ* was 70 ± 6 sec, which is close to the recently determined half-life of 60 s (Scheuer et al. 2022), showing the reliability of our measurement. Ten minutes after Tc addition, the half-life of the *metZ* transcript was increased approximately 5-fold (Table 1). For all analyzed genes including *moeA*, which was strongly affected by the polar effect (Fig. 1B), we detected an increase in the mRNA stability upon Tc exposure (Table 1). Overall, our data suggests that in the presence of Tc, mRNAs are stabilized, and that this contributes to the observed increase in the mRNA levels, unless counteracted by the polar effect.

**Table 1.**
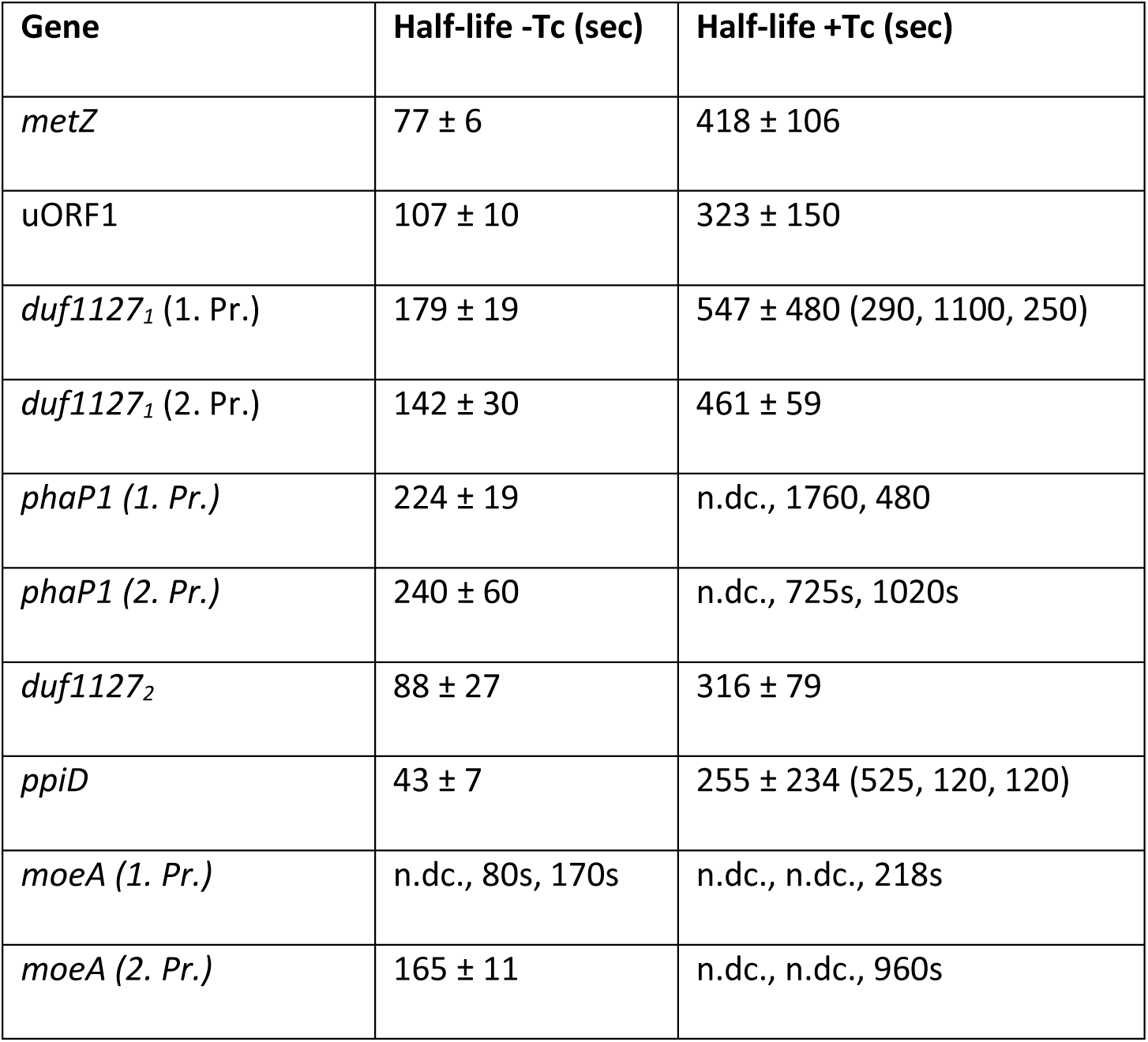
Half-lives of *Sinorhizobium meliloti* mRNA in the absence of Tc (-Tc) and 10 min after addition of 1.5 µg/ml Tc (+Tc) to the cultures. For *duf1127_1_*, *phaP1*, and *moeA*, two primer pairs (1. Pr. and 2.Pr.) were used for the qRT-PCR analysis. N.dc.: no decay was observed.

For the *duf1127_1_* operon, similar, approximately 3-fold increase in the stability was detected using primers targeting the mRNA leader (and thus detecting the uORF-containing sRNA and the uORF-*duf1127_1_* co-transcript) or the *duf1127_1_* sequence (Table 1). Together with the above qRT-PCR results, this suggests that the increase in the *duf1127_1_* mRNA level is not only due to transcript stabilization, but also due to relieved transcription attenuation.

### Strong induction of duf1127_1_ is mediated by its mRNA leader

To address the specificity of the observed mRNA increase upon Tc exposure, additional stresses such as subinhibitory amount of the translational inhibitor chloramphenicol (Cm), H_2_O_2_ (oxidative stress), and heat (42°C) were applied for 10 min. A qRT-PCR analysis of *duf1127_1_* (and its mRNA leader), *phaP1,* and *duf1127_2_* revealed that all three genes were induced by Cm to a similar extend as by Tc (Fig. S3). Although to a less extend, *phaP1* and *duf1127_2_* were also induced by oxidative stress, and *phaP1* showed weak induction by heat (Fig. S3). Thus, only *duf1127_1_* was specifically induced by the translation inhibitors.

To get more insight into the mechanisms of induction by Tc, promoter fusions and translational fusions of *duf1127_1_*, *phaP1,* and *duf1127_2_* were constructed using the *egfp* reporter (Fig. S4). First, reporter expression was analyzed by fluorescence measurement in the absence of Tc. Cultures with the *duf1127_1_* promoter fusion showed high fluorescence, while the fluorescence caused by the translational fusion was very low (Fig. 3A). This suggests that in the absence of Tc, *duf1127_1_* expression is downregulated despite strong promoter activity, and that the 5’-leader is responsible for this downregulation. For *phaP1* and *duf1127_2_*, cultures with the promoter fusions showed fluorescence suggesting moderate promoter activities. The corresponding translational fusions caused approximately 6-fold higher (*phaP1*) and 3-fold higher (*duf1127_2_*) fluorescence. Thus, only *duf1127_1_* is downregulated by its mRNA leader in the absence of Tc (Fig. 3A).

**Figure 3.**
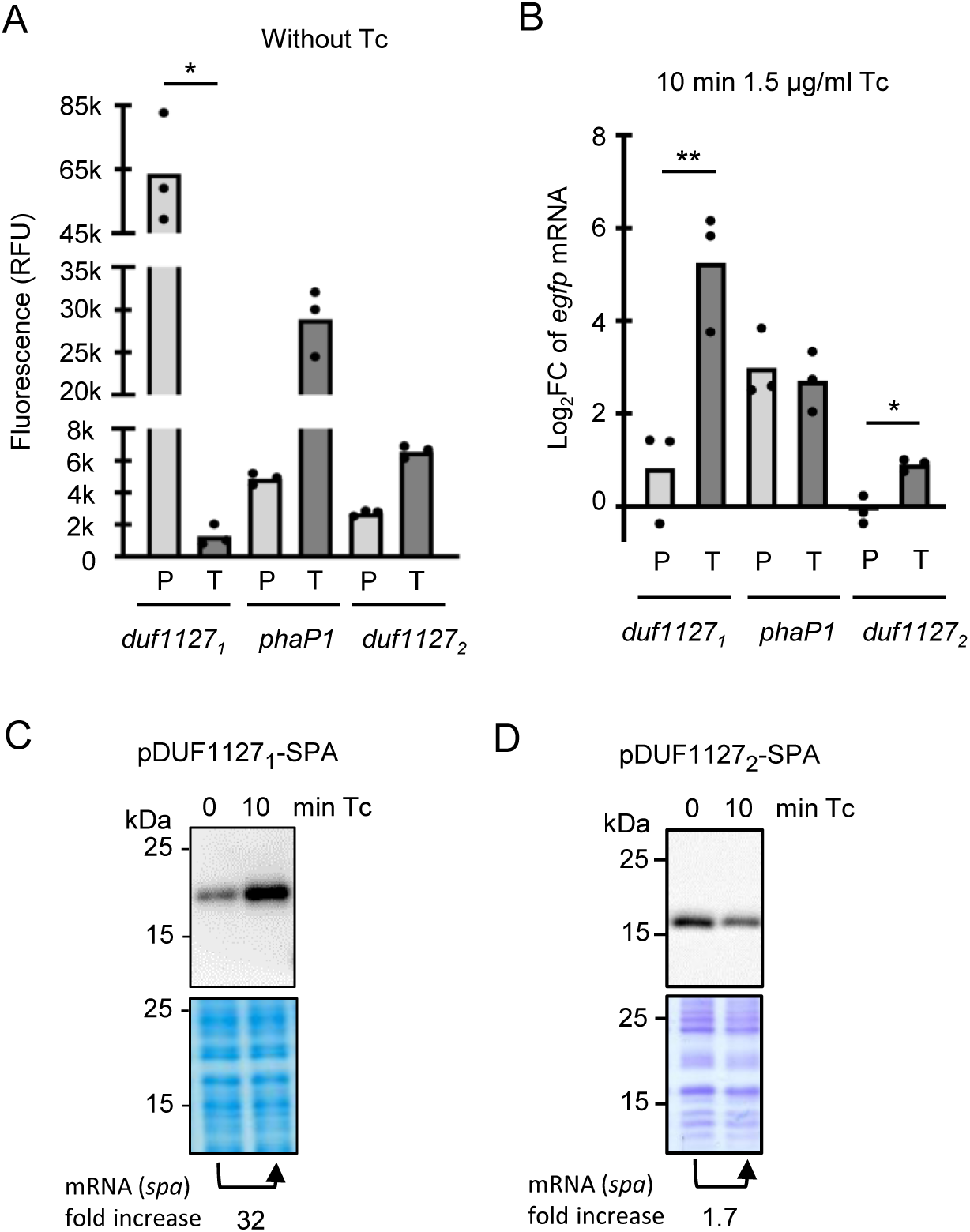
The *duf1127_1_* gene is regulated by its mRNA leader and its induction by tetracycline is detectable at the mRNA and protein level. A) Relative fluorescence of *Sinorhizobium meliloti* cultures in the absence of tetracycline (Tc). The strains harbored *egfp* promoter fusions (P) or translational fusions (T) of the indicated genes on plasmids (Fig. S4). **B)** Analysis by qRT-PCR of changes in the level of the reporter mRNAs upon Tc exposure. Primers targeting *egfp* were used. For other description see A. All graphs show means and single data points of three independent experiments. *** p ≤ 0.001, ** p ≤ 0.01, * p ≤ 0.05. **C) and D)** Western blot analyses with FLAG-directed antibodies of DUF1127_1_-SPA and DUF1127_1_-SPA fusion proteins in cultures before and after Tc addition. The used plasmids are shown in Fig. S4. The bottom panels show Coomassie stained gels with protein probes from the lysates used for the western blots (loading controls). Shown are results of representative experiments. RNA of the same cultures was isolated and analyzed by qRT-PCR with primers directed to the SPA-coding sequence. The fold changes in the mRNA level upon Tc exposure is given below the panels.

Next, using these reporter constructs, changes in mRNA levels 10 min after Tc addition to the cultures were measured by qRT-PCR using *egfp* specific primers (upon such short exposure, fluorescence was not changed). The two *phaP1* reporter fusions showed similar mRNA increase and for *duf1127_2_*, only the translational fusion showed a weak increase (Fig. 3B). Importantly, while the mRNA level of the *duf1127_1_* promoter fusion was not significantly increased, strong increase was observed for the translational fusion (Fig. 3B). We conclude that the leader of *duf1127_1_* mRNA is important for the strong induction by Tc.

To test whether increased mRNA level also leads to increased protein level upon 10 min of Tc exposure, we constructed translational fusions of full-length DUF1127_1_ and DUF1127_2_ to a short SPA-tag (8 kDa; (Babu et al. 2012)), which can be detected with FLAG-directed antibodies. For the *duf1127_1_* construct, increase in the protein level was detected (Fig. 3C). In the same culture, the level of the reporter mRNA was increased 32-fold (primers directed to a sequence encoding the SPA-tag were used). By contrast, the reporter mRNA of the *duf1127_2_* fusion was only slightly increased (1.7-fold), and the amount of the fusion protein was decreased (Fig. 3D).

Together, the results show that upon Tc exposure, *duf1127_1_* is induced at the mRNA and protein level. Overall, the data suggest that the mRNA leader of *duf1127_1_* is involved in transcription attenuation, which is relieved in the presence of Tc .

### Premature transcriptional termination between the uORF and duf1127_1_

To analyze the attenuation mechanism of *duf1127_1_*, we could not directly use plasmid pDUF1127_1_-SPA harboring the abovementioned translational fusion of full-length DUF1127_1_, because mutations accumulated in the construct around the uORF stop codon. Such mutations were not detected when only the first 10 codons of *duf1127_1_* were fused to the tag-coding sequence in a plasmid pDUF^’^-SPA (Fig. 4 and Fig. 5A), and this reporter showed similar regulation in response to Tc (see slopes and lanes labeled with DUF’-SPA in Fig. 5B to 5D). Therefore pDUF^’^-SPA and its derivatives were used in the following experiments.

**Figure 4.**
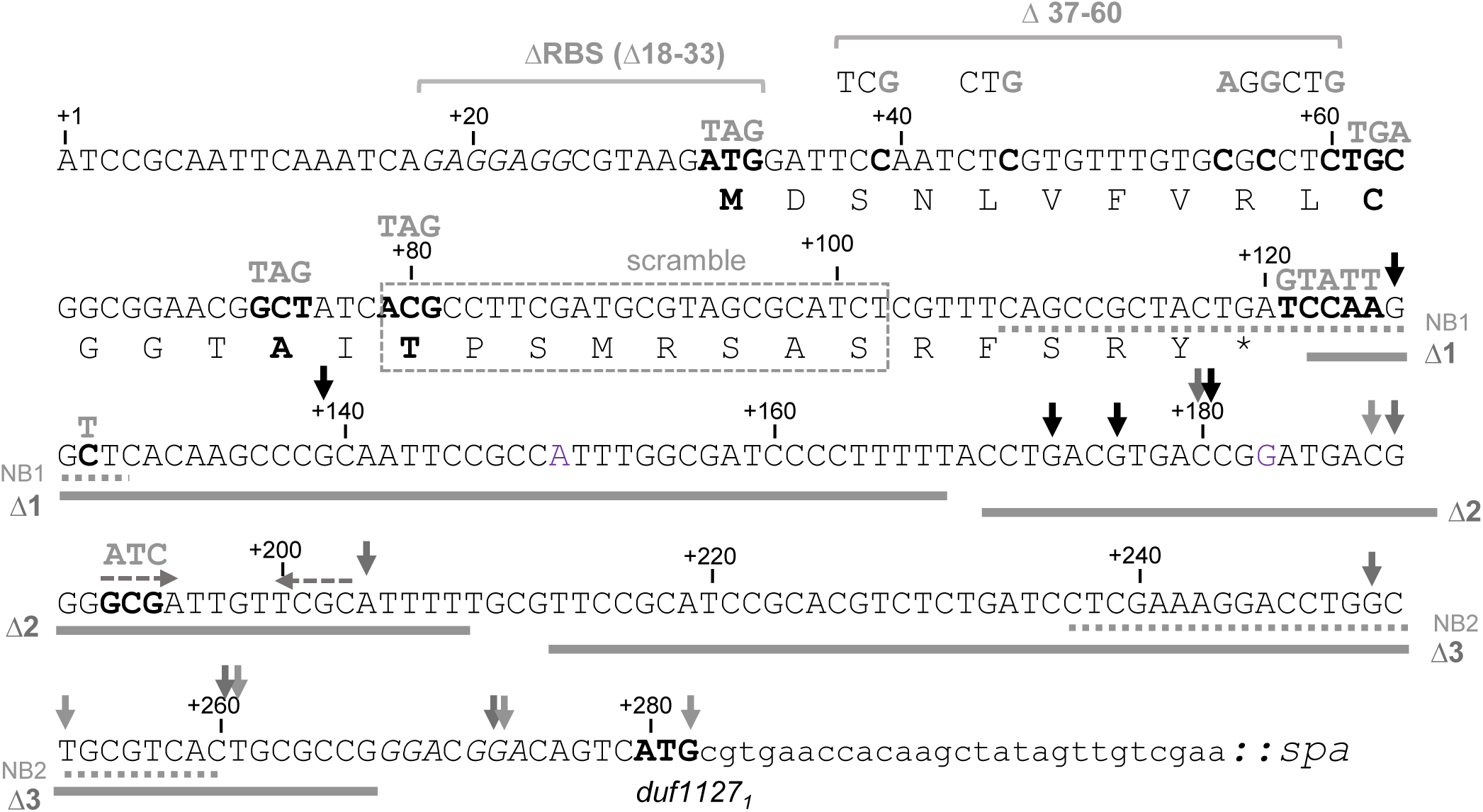
Sequence of the uORF-containing mRNA leader and the first 10 codons of the *duf1127_1_* gene that were used in the translational fusion in the pDUF’-SPA plasmid. +1: Transcriptional start site. The small protein encoded by the uORF is given. Shine-Dalgarno sequences upstream of the uORF and the *duf1127_1_* gene are in italics and the ATG start codons in bold. Mutated nucleotides or codons are also in bold and the corresponding mutations are given above them. Other mutations such as deletions or scrambling are indicated. Sequences targeted by the Northern blot probes (see Fig. 2A and 2B) are underlined with dashed lines. Small vertical arrows mark 3’-ends detected by 3’-RACE.

**Figure 5.**
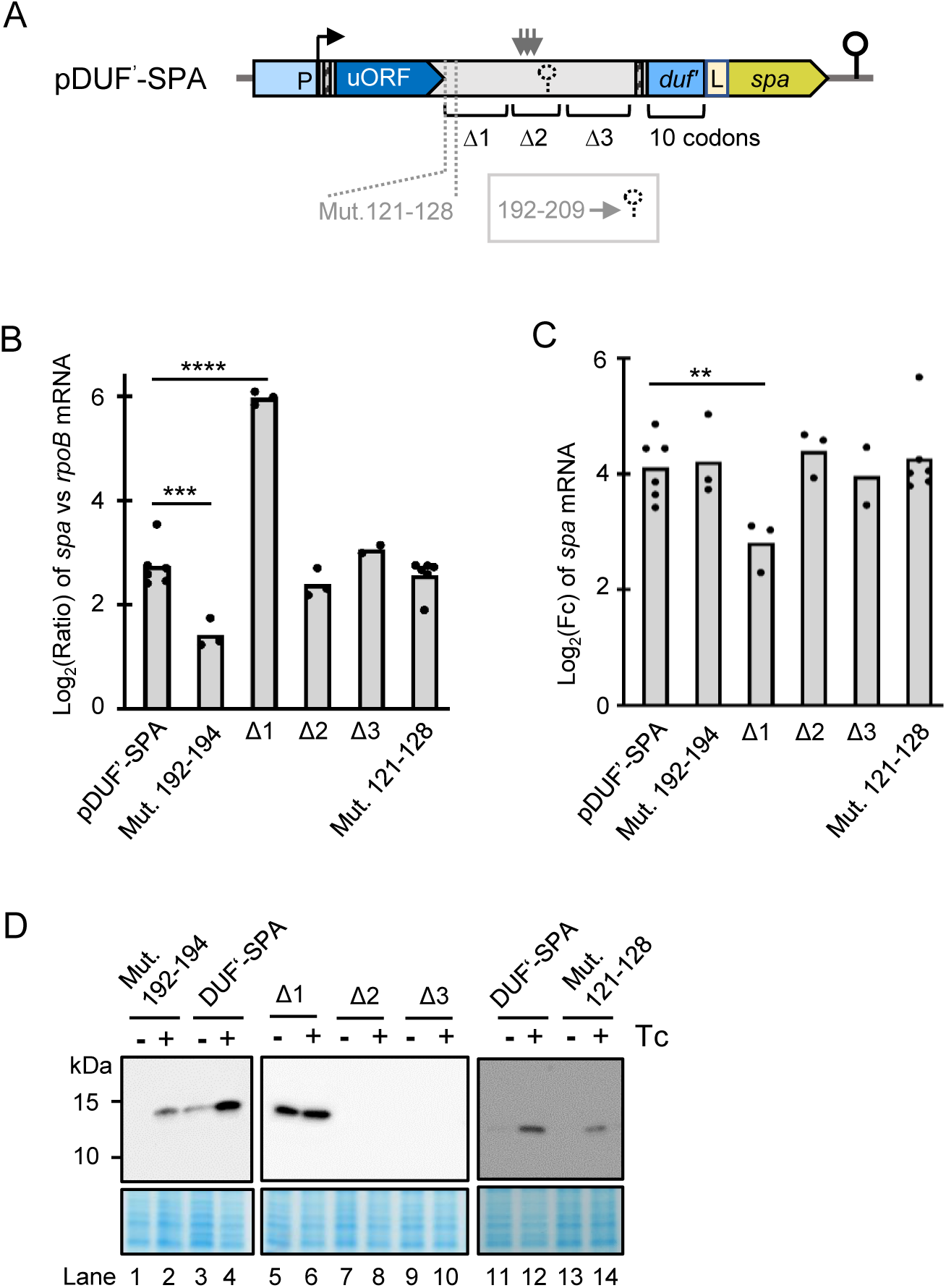
Identification of an attenuation element downstream of the uORF. **A)** Scheme of plasmid pDUF’-SPA. The translational SPA fusion is under the control of the native promotor (P). Flexed arrow: Transcriptional start site. Narrow hatched boxes: Shine-Dalgarno sequences upstream of the uORF and the *duf1127_1_* gene. L: Linker between the in-frame fusion of the first ten *duf1127_1_* codons and the SPA-coding sequence. Indicated are the putative stem-loop structure at transcript positions 192-209 (dashed hairpin), the mutated nucleotides 121-128, the regions Δ1, Δ2 and Δ3 in the intergenic region between the uORF and *duf1127_1_*, and the transcriptional terminator downstream of the cloned fusion (solid hairpin). The detected cluster of 3’-ends downstream of region Δ1 is indicated by vertical arrows. **B)** The qRT-PCR analysis of cultures harboring pDUF’-SPA or indicated derivatives (see A) and grown in the absence of tetracycline (Tc) revealed an attenuation element in the Δ1 region. Primers directed to the SPA-encoding region were used. Calculated was the log_2_ of the ratio between the SPA-encoding transcript and *rpoB* mRNA. Only the Δ1 derivative showed significantly increased basal level of the reporter mRNA. **C)** qRT-PCR analysis of changes in the reporter mRNA levels upon Tc exposure of the cultures used in B. Only the Δ1 derivative showed significantly weaker increase in the mRNA level. **D)** Western blot analyses with FLAG-directed antibodies of the cultures used in C) confirms de-repression of the Δ1 derivative. For other details see Fig. 3C and 3D. All graphs show means and single data points of at least three independent experiments. *** p ≤ 0.001, ** p ≤ 0.01, * p ≤ 0.05.

We first tested whether a Rho-independent terminator is present in the 279 nt mRNA leader of *duf1127_1_*. The only candidate was the sequence <u>GCGA</u>TTGT<u>TCGC</u>ATTTTT (position 192-209 from the transcriptional start site; dyad symmetry regions are underlined; see Fig. 4). To prevent stem-loop formation, GCG in the dyad symmetry was mutated to ATC. According to the qRT-PCR results, this mutation did not increase the *duf’::spa* expression in the absence of Tc and did not affect the induction by Tc. Using Western blot, induction at the protein level was also detected (compare DUF^’^-SPA to Mut. 192-194 in Fig. 5B to 5D). Thus, nucleotides 192-209 do not form a transcriptional terminator.

To identify transcript parts needed for the attenuation, we divided the intergenic sequence between the uORF and *duf’::spa* in three regions, which were deleted separately (Δ1 to Δ3, Fig. 4 and Fig. 5A). The Δ1 deletion led to increased *duf’::spa* expression at the level of mRNA and protein in the absence of Tc (Fig. 5B and 5C), and to weaker induction upon Tc exposure (Fig. 5D). By contrast, the Δ2 and Δ3 deletions did not change the regulation at the level of RNA (Fig. 5B and 5C) and did not lead to strong protein accumulation as observed for Δ1 (Fig. 5D). Thus, the Δ1 region harbors an element that is important for the transcription attenuation upstream of *duf1127_1_*.

Since the Δ1 region starts just downstream of the uORF1 stop codon (Fig. 4), its 5’-part is covered by the ribosome during translation termination under normal conditions. Since in this ribosome-covered part mutations accumulated when pDUF1127_1_-SPA was used, we wondered whether these mutations modulate the expression of the reporter fusion. To test this, we introduced the observed mutations in pDUF’-SPA: transcript nucleotides (nt) 121-128 were mutated from TCCAAGGC to GTATTGGT. However, this sequence change did not affect the *duf’::spa* regulation (Fig.5B to 5D, see slopes and lanes labeled with Mut. 121-128).

Next, we applied 3’-RACE to detect 3’-ends between the uORF and *duf1127_1_* in *S. meliloti* 2011 cultures growing without Tc. The detected 3’-ends are marked with vertical arrows in Fig. 4. A clustering of 3’-ends was observed just downstream of the Δ1 region, between nt 170 and 190 of the transcript (Fig. 4; also marked with vertical arrows in Fig. 5A).

Altogether, the above results show that the Δ1 region (downstream of nt 128 and thus independently of direct ribosomal coverage) is important for the transcription attenuation. They also suggest that the premature transcriptional termination takes place at multiple positions downstream of the Δ1 region.

### Translation of the first half of the uORF is needed to downregulate *duf1127_1_*

Next, we studied the role of the uORF in the regulation of *duf1127_1_*. To analyze the uORF expression, an uORF-SPA translational fusion was constructed (see p-uORF-SPA in Fig. 6A). To address the role of the uORF, in plasmid pDUF^’^-SPA, its start codon was mutated to a stop codon (M1/Stop mutation destroying the uORF; Fig. 6A).

**Figure 6.**
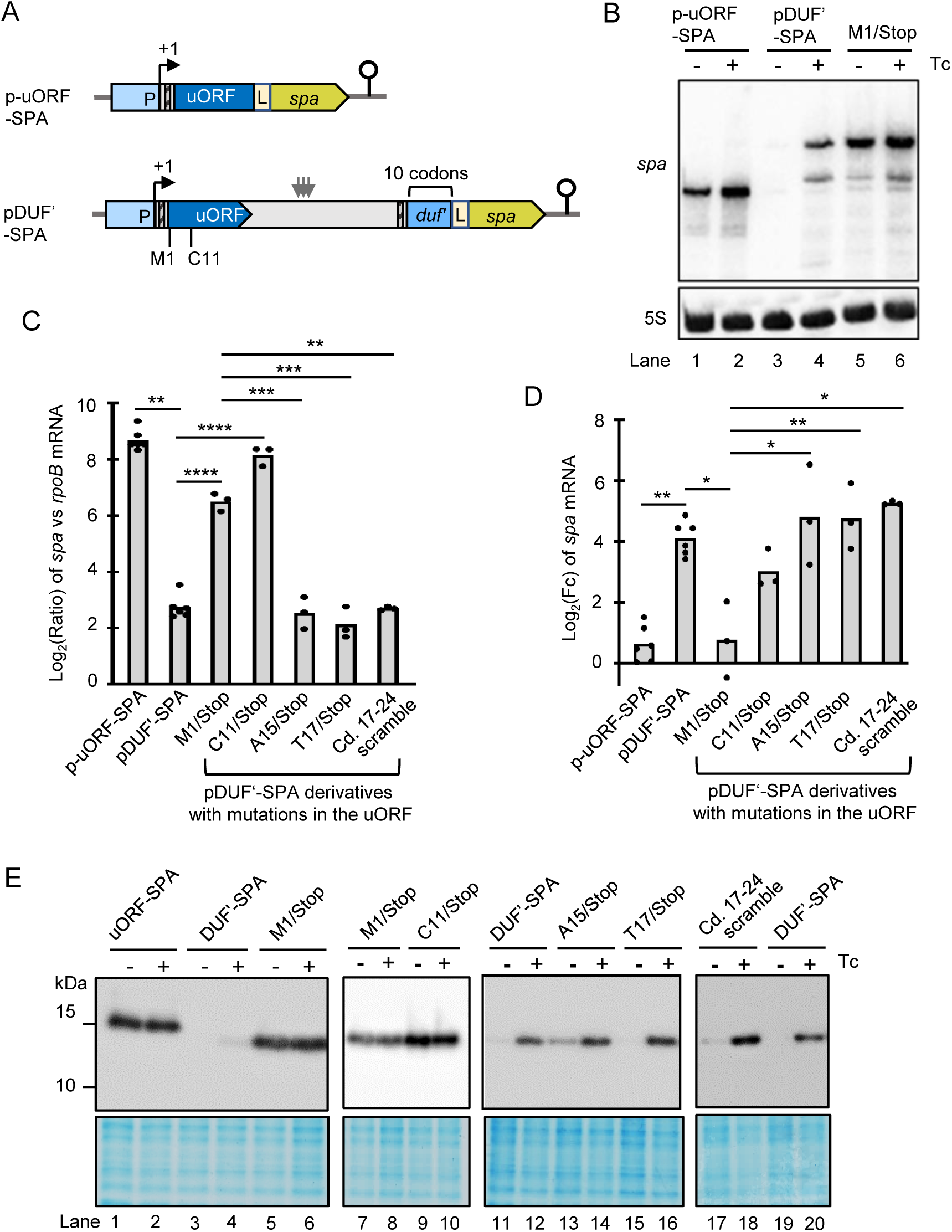
Translation of the first half of the uORF enables regulation of *duf1127_1_*. **A)** Schematic representation of p-uORF-SPA und pDUF’-SPA. In the latter, uORF codons, the mutation of which to stop codons led to relieve of attenuation, are indicated. For other details see Fig. 5A. **B**) Northern blot analysis of cultures harboring p-uORF-SPA, pDUF’-SPA or the M1/Stop derivative of pDUF’-SPA. RNA was isolated 10 min after addition of tetracycline (+Tc) or its solvent ethanol (-Tc). The membrane was hybridized with a probe directed to the SPA-encoding sequence and rehybridized with a probe against 5S rRNA (loading control). **C)** qRT-PCR analysis of cultures harboring the indicated plasmids and grown in the absence of Tc. Calculated was the log_2_ of the ratio between the SPA-encoding transcript and *rpoB* mRNA. For other description see Fig. 5B. The results show the basal expression of the reporter mRNA from the different plasmids, indicating constitutive uORF expression and de-repression of *duf’::spa* due to the M1/Stop and C11/Stop mutations. **D)** qRT-PCR analysis of changes in the reporter mRNA levels upon Tc exposure of the cultures used in C). The results suggest constitutive uORF1 expression and no significant mRNA increase (induction) of the de-repressed M1/Stop transcripts. The C11/Stop transcript, albeit strongly accumulated in the absence of Tc, showed mRNA increase, which was not significantly different from that of the induced pDUF’-SPA and the de-repressed M1/Stop. Induction of A15/Stop, T17/Stop and the scramble mutant was similar to the parental construct and significantly different from M1/Stop. **E**) Western blot analyses with FLAG-directed antibodies of the cultures used in D confirms constitutive expression of the uORF and de-repression of M1/Stop and C11/Stop. All graphs show means and single data points of at least three independent experiments. *** p ≤ 0.001, ** p ≤ 0.01, * p ≤ 0.05.

When p-uORF-SPA was used, the reporter mRNA was detected by Northern hybridization in cultures without Tc (Fig. 6B). Strong expression at the level of RNA in the absence of Tc was confirmed by qRT-PCR (Fig. 6C). The Northern blot and qRT-PCR analyses revealed a slight increase in the level of this mRNA upon Tc exposure (Fig. 6B and 6D). Using Western blot, the corresponding uORF1-SPA fusion protein was detected in high and similar amounts in cultures with and without Tc (Fig. 6E, compare lanes 1 and 2 showing uORF-SPA to lanes 3 and 4 supposed to show the Tc induction of the DUF’-SPA control, which is barely visible because of the short detection time). Thus, the uORF is efficiently transcribed and translated in the absence of Tc.

In line with results shown in Fig. 2 and Fig. 5 above, when pDUF^’^-SPA was used, the reporter mRNA *duf’::spa* was detectable by Northern hybridization only after Tc addition and strong mRNA increase was detected by qRT-PCR (Fig. 6B to 6D). Importantly, when the uORF was destroyed by the M1/Stop mutation, the *duf’::spa* expression was de-repressed in cultures without Tc. This de-repression (lack of attenuation) was detected at the levels of mRNA (Fig. 6B and 6C) and protein (lanes 5 and 6 in Fig. 6D). Thus, in the absence of Tc, translation of the uORF is necessary for attenuation of *duf1127_1_*.

Next, we asked whether a specific part of the uORF must be translated for regulation of *duf1127_1_*. We constructed pDUF^’^-SPA derivatives, in which one of the codons C11, A15, or T17 in uORF was mutated to a stop codon, and respective cultures were analyzed by qRT-PCR and Western blot (Fig. 6C to 6E). In the absence of Tc, the *duf’::spa* expression from the C11Stop construct was strong (de-repressed) at the level of RNA (Fig. 6C) and protein (lanes 9 and 10 in Fig. 6E). Upon Tc exposure, this construct still led to increase in the mRNA level. The increase was weaker than that of the parental construct pDUF^’^-SPA, but the difference was not statistically significant. However, there was also no significant difference in induction when compared to the de-repressed construct M1/Stop (Fig. 6D). In contrast, when the uORF translation was terminated at codon 15 or 17, attenuation in the absence of Tc and induction upon Tc exposure were similar to those of the parental construct and statistically different from M1/Stop (Fig. 6C and 6D; lanes 11 to 16 in Fig. 6E). This shows that the first half of the uORF must be translated for a proper *duf1127_1_* regulation.

Most of the described mechanisms for ribosome-dependent transcription attenuation rely on RNA sequence elements in the nascent RNA that can undergo alternative base-pairing or change their accessibility for the transcription factor Rho (Turnbough 2019). Knowing that the bacterial ribosome covers approximately 30 nt (Yusupova et al. 2001); (Hadjeras et al. 2023), we first reasoned that ribosome occupancy of the uORF codons 17 to 20 could be important for the transcription attenuation of *duf1127_1_*. In the absence of Tc, these codons would be covered by the ribosome terminating translation at A15/Stop, but would be accessible if the ribosome terminates translation at C11/Stop. However, scrambling the uORF codons 17 to 24 (nt 78 to 102 of the transcript, see Fig. 4) did not affect the regulation of the *duf’::spa* reporter at the level of RNA (Fig. 6C and 6D) and protein (lanes 17 to 20 in Fig. 6E). Since the scrambling drastically changed the sequence and possible RNA structures, we conclude that the second half of the uORF is not involved in the regulation of *duf1127_1_*.

### The first third of the uORF contains an attenuation element that can be occluded by a ribosome or masked by an RNA structure

Based on the above results, we proposed that an uORF sequence, which is translated in the A15/stop reporter construct, but is not covered by the ribosome during translation termination at the 15^th^ codon, contains an attenuation element. According to this hypothesis, for successful attenuation the first 10 codons of the uORF (nt 31 to 60 of the transcript, see Fig. 4) must be accessible for a factor such as Rho, and this accessibility depends on their translation. In this scenario, in the absence of Tc, nucleotides 31 to 60 would be transiently free when a ribosome passes them and the next ribosome is still not bound to the ribosome binding site (RBS) of the uORF. Upon Tc exposure, nt 31 to 47 (the first 6 uORF codons) would be blocked by a ribosome with impaired translation initiation, thus rendering the proposed attenuation element partly inaccessible. On the other hand, in the case of the C11/Stop construct in the absence of Tc, the ribosome will block nt 47 to 60 (codons 6 to 10) during the process of translation termination at the 11^th^ codon, and the proposed attenuation element will also be partly blocked. This scenario explains the Tc-induction of the parental *duf’::spa* construct and of its A15/Stop or T17/Stop derivatives, as well as the de-repression of its C11/Stop derivative in the absence of Tc.

We considered that the putative attenuation element could be a Rho-binding sequence harboring C > G. Therefore, in pDUF’-SPA the C-rich codons 3, 5, 9 and 10 were replaced by synonymous codons, thus raising the G and lowering the C content of this uORF region (Fig. 4, Fig. 7A). In line with our expectations, these mutations led to de-repression of *duf’::spa* in the absence of Tc (Fig. 7B and 7C; lanes 1 and 2 in Fig. 7D), showing that the RNA and not the peptide sequence is important for the observed regulation. Deletion of uORF codons 3 to 10 in pDUF’-SPA (Δ37-60) had similar effect: de-repression in the absence of Tc at the level of RNA (Fig. 7B) and protein (lanes 7 and 8 Fig. 7D) and very weak RNA increase upon Tc exposure (Fig. 7C).

**Figure 7.**
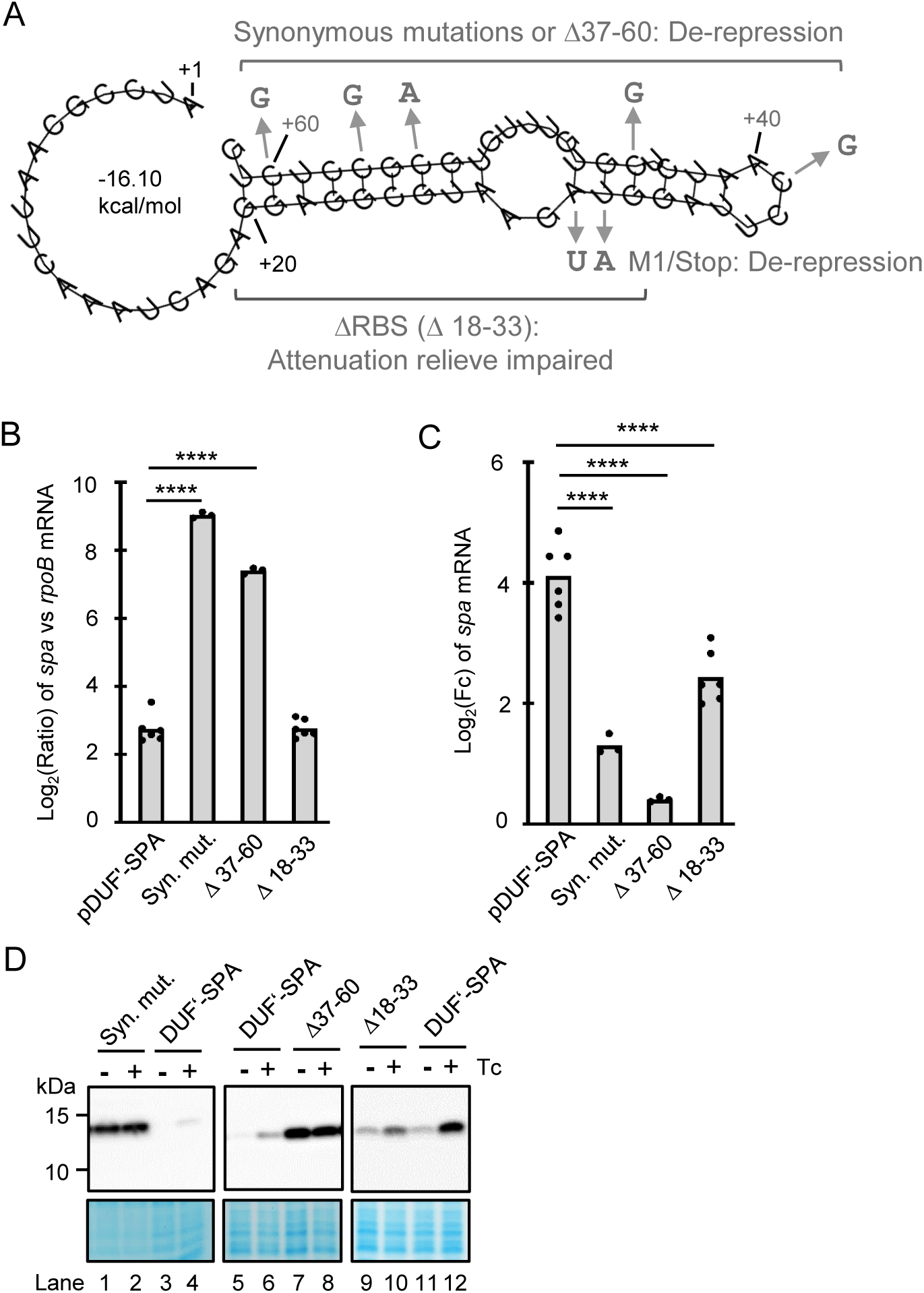
An attenuation element in the first third of the uORF1 is accessible upon unhindered translation. **A)** Predicted secondary structure of the first 62 nt nucleotides of the *duf1127_1_* mRNA leader. Indicated are mutations causing attenuation relieve (de-repression) or relieve impairment in pDUF’-SPA derivatives. **B)** qRT-PCR analysis of cultures harboring pDUF’-SPA or its indicated derivatives and grown in the absence of Tc shows relieved attenuation upon synonymous mutations in the transcript region 37-60 or a deletion of this region (see panel A and Fig. 4). Calculated was the log_2_ of the ratio between the SPA-encoding transcript and *rpoB* mRNA. For other description see Fig. 5B. **C)** qRT-PCR analysis of changes in the reporter mRNA levels upon Tc exposure of the cultures used in C. All three pDUF’-SPA derivatives showed significantly lower increase in the reporter mRNA amount compared to the parental construct. **D)** Western blot analyses with FLAG-directed antibodies of the cultures used in D confirms relieve of attenuation due to the synonymous mutations in the transcript region 37-60 or the deletion of this region. Additionally, it confirms the impaired relieve of attenuation by Tc for the Δ18-33 derivative that lacks the ribosome binding site of the uORF.

However, the above hypothesis cannot explain the de-repression of *duf’::spa* in the M1/Stop construct, in which the uORF is not translated because it lacks a start codon and thus a functional RBS. This de-repression could be explained by a secondary structure, which blocks the attenuation element. Indeed, using RNAfold we predicted that the sequence comprising the RBS (the Shine-Dalgarno sequence and the start codon) and the first 10 codons of the uORF can fold in a hairpin structure (Fig. 7A), and a similar structure was predicted for the M1/Stop derivative (Fig. S5). Thus, a deletion of the RBS of the uORF is expected to constitutively expose the C-rich attenuation element. We tested this by deleting nt 18 to 33 of the transcript (Fig. 7A). In the absence of Tc, the Δ18-32 construct showed low reporter mRNA levels, which indicated transcription attenuation (compare Δ18-33 and pDUF’-SPA in Fig. 7B), although the uORF could not be translated. This is in strong contrast to the de-repressed M1/Stop construct (see Fig. 6 above). These results support the existence of an RNA structure that masks the attenuation element in the uORF. Moreover, the results imply that under conditions of no timely ribosome binding to the nascent RNA, this RNA structure is formed and *duf1127_1_* is induced even in the absence of antibiotics.

Upon Tc exposure, the Δ18-33 construct showed significantly lower increase in the level of the reporter mRNA compared to pDUF’-SPA (Fig. 7C). The Western blot result also suggested weaker induction at the protein level for the Δ18-33 construct compared to pDUF’-SPA (Lanes 9 to 12 in Fig. 7D). The increase in the mRNA level of the Δ18-33 construct with constitutively exposed attenuation element was most probably due to the increased mRNA stability in the presence of Tc. Of note, the Δ18-33 construct was the only construct tested in this work, which showed impaired mRNA induction upon Tc exposure although its attenuation in the absence of Tc was normal. All other constructs with impaired Tc-induction were de-repressed in the absence of Tc (see Δ1 in Fig. 5; M1/Stop in Fig. 6; syn. mut. and Δ37-60 in Fig. 7). This suggests that in the Δ18-33 construct, the attenuation element encompassing nt 37-60 is permanently exposed and transcriptional attenuation does not take place. The increased level of *duf’::spa* mRNA of the Δ18-33 construct upon Tc exposure could be explained by mRNA stabilization. Altogether, the results support the existence of an attenuation element in the first third of the uORF, which can be masked by a ribosome stalled at the begin of the uORF and by an RNA structure under conditions of no timely ribosome binding to the nascent mRNA.

## Discussion

Our RNA-seq analysis revealed large transcriptome changes 10 min after addition of a subinhibitory Tc amount to *S. meliloti* 2011 cultures. The data suggest that generally, the effects are due to co- and post-transcriptional mechanisms such as PTT and RNA stabilization, which caused polar effects in operons. Furthermore, we revealed a fast and strong response to Tc exposure of the *duf1127_1_* gene, which is mediated by a 5’-UTR that senses translation efficiency in the cell.

Most available studies on transcriptome changes in bacteria exposed to antibiotics reflect exposure times longer than 10 min (typically from 30 min to several hours) and report on changes of hundreds of genes grouped in GO and KEGG terms (Zaide et al. 2023); (Cianciulli Sesso et al. 2021). It is expected that, among expression changes directly supporting the adaptation to antibiotic stress, long exposure times cause secondary effects. Despite this limitation, it is noteworthy that exposure to translation inhibitors was repeatedly found to upregulate genes involved in translation, including ribosomal genes (Kesavan et al. 2020); (Cianciulli Sesso et al. 2021)). In our RNA-seq analysis, “structural constituent of ribosome” was the only significantly enriched GO term of upregulated genes 10 min after addition of Tc (0.75 of the minimal inhibitory concentration(MIC)) to *S. meliloti* cultures. We were surprised to detect expression changes in hundreds of genes (Table S1), since, in comparison, in a recent study analyzing *A. baumannii* after 10 min exposure to 0.5-MIC of minocycline, only 25 DEGs were detected (Gao und Ma 2022). The lower antibiotic concentration used by Gao et al. is probably a reason for the low number of significantly affected genes. We also detected weaker mRNA changes for selected genes of *S. meliloti* when using lower Tc concentrations (Fig. S2).

In our study, the principal lack of functional groups (besides ribosomal genes) among *S. meliloti* DEGs suggests that general mechanisms acting on RNA-level were influenced by Tc-caused ribosome stalling, particularly at start codons. The observed polar effects in operons (RNA increase at the begin and decrease toward the end of polycistronic transcripts; Fig. 1B; Table S1-a and Table S1-b), probably result from two mechanisms with opposing effects. On the one hand, the increase in the mRNA level of first genes in transcripts could be explained by mRNA stabilization, since ribosomes trapped by Tc at the stage of translation initiation probably block RNA degradation by the major RNases RNase E and RNase J, which are known to have 5’ to 3’ polarities (Hui et al. 2014). The decrease in the mRNA level of downstream genes in polycistronic transcripts probably reflects the decoupling between translation and transcription, which leads to PTT in gram-negative bacteria (Zhu et al. 2019); (Johnson et al. 2020). Toward 3’-ends of polycistonic operons, the PPT effect is obviously stronger than the effect of mRNA stabilization, since *moeA* mRNA was stabilized, but its levels was significantly decreased upon Tc exposure (Table 1, Fig. 1B).

We did not analyze the mRNA stability changes transcriptome-wide, but detected transcript stabilization for all tested genes regardless whether they showed mRNA increase, decrease or no change upon Tc exposure (Table 1). This suggests that mRNA stabilization is a general effect under translation inhibiting conditions. Stabilization of mRNA upon translation inhibition was mentioned previously for *cspA* mRNA in *E. coli* treated with chloramphenicol (Jiang et al. 1993). The here determined mRNA half-lives in the absence of Tc are very short: of the seven tested transcripts, four showed half-lives below 2 min, which is in line with a recent study showing that bacterial mRNA is even more unstable than generally assumed (Jenniches et al. 2024). The most unstable mRNAs were *metZ* and *ppiD*, with half-lives of 77s and 43s, respectively. Both genes are first in operons and their mRNA was stabilized in the Tc-samples 3- to 5-fold (Table 1). However, according to RNA-seq, *metZ* was not deregulated and *ppiD* had an increase below the used threshold (Table S1). This could be explained by the length of these genes (*metZ* :1185 nt and *ppiD*: 1892 nt), which probably led to PTT toward their 3’-ends.

Our data suggest that mRNA stabilization and PTT together shaped the *S. meliloti* transcriptome shortly after addition of Tc to the cultures. Thus, position of genes in operons has influence on changes in their transcript abundance under conditions of translation inhibition. Previously, position of genes in bacterial operons was also shown to influence their mRNA stabilities, with mRNA segments downstream in polycistronic transcripts being more stable (Selinger et al. 2003); (Steglich et al. 2010). Thus, positions of genes in operons and length of operons are probably under evolutionary pressure for optimization of gene expression under changing environmental conditions. Along this line, some iTSS, known to be often present in bacterial operons (Sharma et al. 2010), (Schlüter et al. 2013), (Čuklina et al. 2016), might evolved to counteract positional effects.

In addition to the abovementioned mechanisms operating transcriptome-wide, individual genes are under specific regulation, a point that we addressed for three genes using promoter and translational reporter fusions. The results suggested that increased promoter activity contributes to *phaP1* (encoding phasin) upregulation upon Tc exposure (Fig. 3A and 3B). Upregulation of *phaP1* under various stresses was observed in this study (Fig. S3) and by others (McIntosh et al. 2021). For *duf1127_2_*, our data suggest that mRNA stabilization is the major factor responsible for upregulation under Tc conditions (Table 1 and Fig. 3), in agreement with its small gene size and monocistronic organization. All six *duf1127* homologs listed in Table S1-b are small and all except *duf1127*_1_ are in monocistronic transcripts, characteristics explaining their enrichment among the upregulated DEGs. While the *duf1127_2_* gene was slightly induced by oxidative stress, *duf1127*_1_, responded specifically to translation inhibition (Fig. S3). According to our data, *duf1127*_1_ induction was due to relieve of transcriptional attenuation in the 5’-UTR, in line with its very fast response to Tc exposure (Fig. 2D).

The uORF1-based transcriptional attenuation of *duf1127*_1_ does not depend on an intrinsic terminator (Fig. 5) and depends therefore probably on the transcriptional termination factor Rho. Involvement of Rho is also supported by the observation that deletion of C-rich sequences (Δ37-60 and Δ1 region, Fig. 4) led to de-repression of the *duf’-spa* fusion in the absence of Tc (Fig. 5 and Fig. 7 ).The importance of the cytidines in the codons 3 to 10 (nt 37-60 if the transcript) was also shown by their mutational replacement (Fig. 7). However, both sequences, designated above “attenuation elements”, are shorter than a typical *rut* site, which usually encompasses 70-90 nt (Fig. 4; (Kriner et al. 2016); (Molodtsov et al. 2023)). Thus, we speculate that the two attenuation elements could represent a discontinuous *rut* site, with a part located in the first third of the uORF and a second part downstream of the uORF.

Based on the presented results, we propose the following model for transcription attenuation of *duf1127*_1_ (Fig. 8): Under optimal growth conditions, two attenuation elements in the nascent RNA ensure Rho-dependent stop of transcription between the uORF and *duf1127*_1_. The first attenuation element is located at codons 3 to 10 of the uORF and is accessible after it is translated by a pioneering ribosome and before it is occupied by a next ribosome (Fig. 6 and Fig. 7). The second element is located downstream of the uORF (Fig. 5). In the presence of Tc, the translation initiation by the pioneering ribosome is impaired. The ribosome stalled at the initiation codon occludes a half of the first attenuation element and therefore attenuation is relieved, leading to transcription of *duf1127_1_.* A similar effect is expected if translation elongation at the first ten uORF codons is impaired. Furthermore, our data suggest that a delay in the occupancy of the uORF RBS by a pioneering ribosome would also result in relieve of attenuation, because the nascent RNA forms a secondary structure that blocks the first attenuation element (Fig. 7). The presented model suggests that the mRNA leader of *duf1127_1_* is well suited to sense translation inhibition, ribosome shortage and shortage of charged tRNAs.

**Figure 8.**
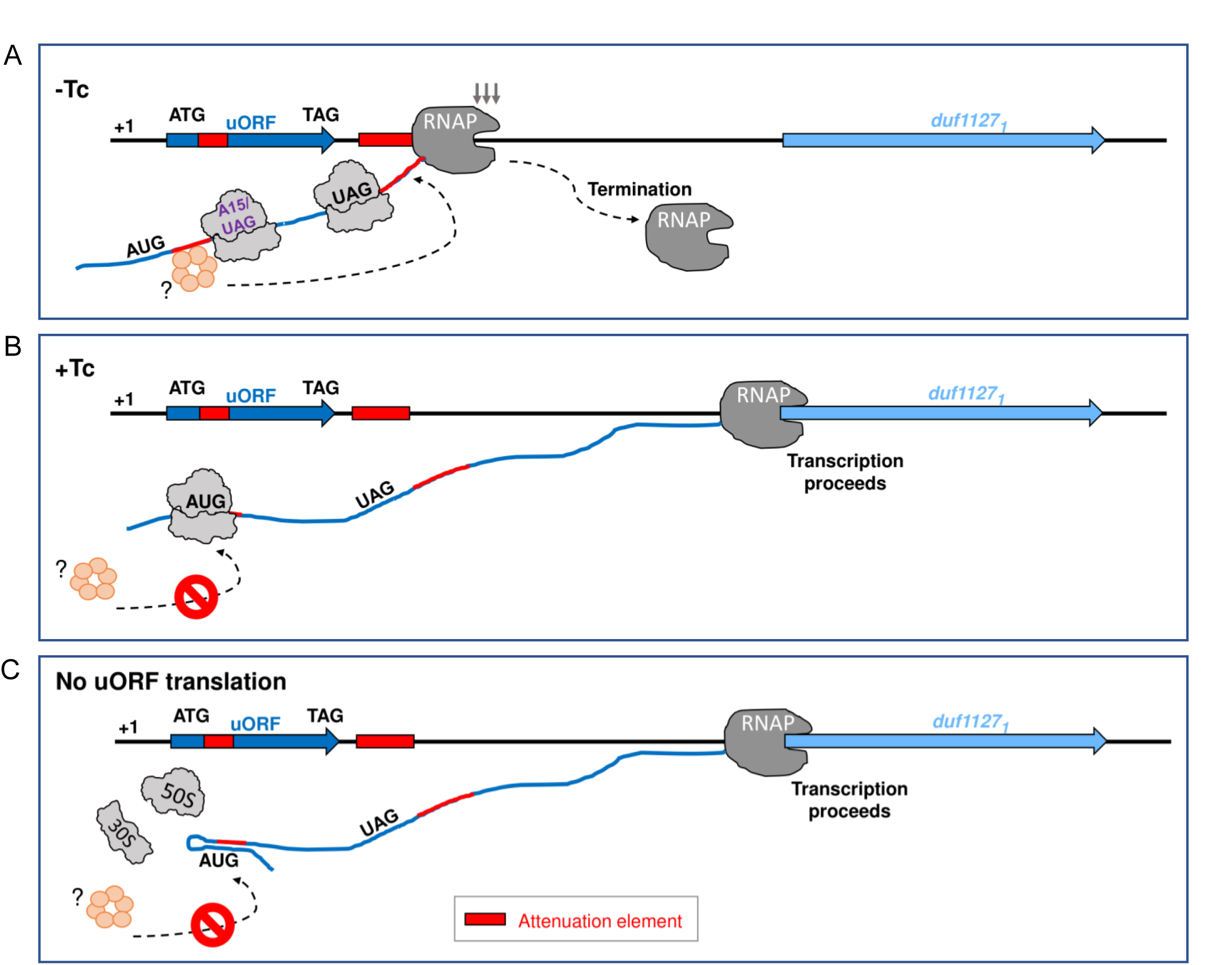
Model of the attenuation mechanism controlling *duf1127_1_* expression. Our data suggest that two sequence elements (marked in red) are needed for transcriptional attenuation of *duf1127_1_* in the absence of tetracycline (Tc). The first element is located at codons 3 to 10 of the uORF and is accessible under conditions of normal translation, while the second element is located downstream of the uORF. **A)** For successful attenuation in the absence of Tc, a ribosome must translate at least the first 14 uORF codons (translation termination at codon 15 still allows for attenuation). In the time window before the next ribosome occupies the ribosome binding site (RBS) of the uORF, the translated attenuation element is accessible for a factor (presumably the the Rho hexamer), which then also binds the second attenuation element in the nascent RNA and leads to premature transcriptional termination. **B)** In the presence of Tc, the translation initiation at the start codon of the uORF is impaired. The stalled ribosome blocks the accessibility to the first attenuation element and therefore transcription proceeds into *duf1127_1_* (attenuation is relieved). **C)** According to our data, a delay in the occupancy of the uORF RBS by a pioneering ribosome would also lead to relieve of attenuation and *duf1127_1_* transcription, because the nascent RNA forms a secondary structure that blocks the first attenuation element.

Rho-dependent transcriptional attenuation regulated by a *rut* site occlusion due to ribosome pausing was described previously. An example is the tryptophanase operon *tnaCAB* in *E. coli* (reviewed in (Turnbough 2019)), where a *rut* site immediately downstream of the uORF *tnaC* (24 aa) is accessible for Rho if the ribosome efficiently terminates translation. Under high cellular tryptophan concentrations, the ribosome is stalled at Pro24 in *tnaC*, thus occluding the *rut* site. Another example is the copper-regulated attenuation of a transporter operon in *Bordetella pertussis* (ref). In this case, translation arrest near the 3’-end of the uORF bp2923 leads to masking of the *rut* site, which is located in the uORF. Perception of copper relieves the translation arrest and the successful uORF translation leads to accessibility of the *rut* site and attenuation of the downstream genes (ref). Similar to *duf1127*_1_, the uORF bp2923-mediated attenuation in *B. pertussis* depends on a translated *rut* site, which is accessible in a time window between two translating ribosomes. In contrast to the *B. pertussis rut* site located at the end of the uORF bp2923, the here described *S. meliloti* attenuation element is located at the begin of the uORF, a position that probably evolved to ensure its immediate masking by an RNA structure under conditions of low ribosome availability.

The functions of *duf1127*_1_ and of the arginine rich DUF1127 proteins in general remain unclear. DUF1127 genes are mostly present in Gamma- and Alphaproteobacteria, the latter often harboring multiple homologs (Kraus et al. 2020). In *Rhodobacter sphaeroides*, a DUF1127 protein was described as an RNA-binding protein regulating stress-related sRNAs (Billenkamp et al. 2015), (Grützner et al. 2021). Another *R. sphaeroides* DUF1127 protein was shown to be transiently induced upon heat shock (McIntosh et al. 2021). In *Agrobacterium tumefaciens* having thee short and three longer DUF1127 genes, deletion of the three short genes led to phenotypic changes at late growth phases and influenced carbon and phosphate metabolism (Kraus et al. 2020). In *Brucella*, the three short DUF1127 genes are linked to fucose utilization (Budnick et al. 2018). The abovementioned, triple *A. tumefaciens* mutant showed increased biofilm formation (Kraus et al. 2020), a phenotype recently reported for a *Vibrio vulnificus* DUF1127 deletion mutant (Feng et al. 2024). The *S. meliloti duf1127*_1_ corresponds to (give the gene ID) in *A. tumefaciens,* one of the large DUF1127 genes with unknown function (Kraus et al. 2020). Our analyses revealed an uORF1-based mechanism that allows for *duf1127*_1_ regulation not only in response to translation inhibition, but also to ribosome availability. This suggests that the 5’-UTR of *duf1127*_1_ senses the translational status in the cell und could be important for the cellular homeostasis in *S. meliloti*.

In summary, we found that the early response of *S. meliloti* to Tc exposure is characterized by posttranscriptional mechanisms and uncovered an unusual, uORF-mediated transcription attenuation mechanism of a DUF1127 gene .

## Material and methods

### Strains cultivation and conjugation

*Sinorhizobium meliloti* 2011 was cultivated in TY (Beringer 1974) with 250 μg/ml streptomycin. Thirty ml cultures were grown in 50 ml Erlenmeyer flask at 30°C and 140 r.p.m. until OD_600nm_ of 0.5. When appropriate, gentamicin (10 µg/ml) was added to maintain plasmids. Tetracycline (Tc) concentrations, which were applied for antibiotic stress, are stated. *Escherichia coli* was cultivated in LB broth. For standard cloning, *E. coli* DH5α was used. Shuttle plasmids were transferred to *S. meliloti* by diparental conjugation using *E. coli* S17-1 (Simon et al. 1983)

### Plasmid construction

The used oligonucleotides (Microsynth, Balgach, Switzerland) are listed in Table S5 and plasmids in Table S6. Cloning was performed as described (Scheuer et al., 2022; , (Sambrook et al. 1989). Enzymes were purchased from Thermo Fischer Scientific. After cloning in plasmid pJET1.2/blunt (Thermo Fischer Scientific), inserts were subcloned into the shuttle plasmid pSW1 (Hadjeras et al. 2023) or pRS1 derivatives (Scheuer et al. 2022) and sequenced with plasmid-specific primers (Microsynth Seqlab, Göttingen, Germany). The reporter sequences were adapted to the *S. meliloti* codon usage (McIntosh et al. 2008).

To generate promoter fusions with *egfp* reporter, approximately 200 bp upstream of the respective transcription start site (TSS) were amplified and cloned upstream of a promoterless *egfp* gene preceded by a Shine-Dalgarno sequence (SD) in pRS1-sSD-egfp (Scheuer et al. 2022). For translational *egfp* fusions, the same respective upstream region harboring the promoter was amplified together with the corresponding mRNA leader and ten codons (or the full-length coding sequence) of the gene of interest and cloned in frame to the third *egfp* codon in pRS-‘egfp (Scheuer et al. 2022). The promoter, leader and coding regions of *duf1127_1_* and *duf1127_2_* were also cloned in frame to a short linker and a SPA-tag encoding sequence in pSW1 (Hadjeras et al. 2023). Site directed mutagenesis was first performed on sequences cloned in pJET and then sub-cloned in the respective shuttle plasmid. To delete specific regions of cloned sequences, the upstream and downstream regions flanking the sequence to be cloned were amplified and then fused using overlapping PCR. Scramble mutations were introduced in oligonucleotides that were used for PCR, followed by overlapping PCR.

### Treatment with Tc and other stressors

Unless stated otherwise, Tc (stock solution of 10 mg/ml in ethanol) was added to the exponentially growing *S. meliloti* cultures to a final concentration of 1.5 µg/ml and cells were harvested 10 min thereafter. To parallelly growing control cultures, the same volume of the solvent ethanol was added. No difference in gene expression was detected between such control cultures and cultures to which solvent was not added. Similarly, *S. meliloti* cultures were treated for 10 min with chloramphenicol (final concentration of 9 µg/ml), 1 mM H_2_O_2_ or were shifted for 10 min to 42°C.

### RNA isolation

Routinely, RNA was isolated from 15 ml *S. meliloti* culture (OD_600 nm_ = 0.5) poured in centrifugation tubes filled with ice rocks and centrifuged at 6000 g and 4°C. The bacterial pellet was resuspended in 1 ml TRIzol® reagent (Thermo Scientific, Waltham, USA) and RNA was isolated according to the manufacturer instructions. Additional hot acidic phenol and chloroform-isoamyl alcohol (24:1) extractions were conducted. RNA was precipitated with ethanol, washed with 75 % ethanol, dried and dissolved in ultrapure water.

To isolate RNA for determination of mRNA half-lives, at the indicated time points after rifampicin addition (final concentration of 600 µg/ml), 500 µl of a bacterial culture was withdrawn and mixed with 1 ml Bacteria Protect Reagent (Qiagen). Then 1 ng of a spike-in *in vitro* transcript (*Sulfolobus solfataricus rrp41*; (Scheuer et al. 2024)) was added and RNA was isolated using RNeasy Mini Kit (Qiagen).

### RNA-seq, identification of DEGs and TSS, and GO and KEGG analyses

RNA-seq was performed by vertis Biotechnologie AG (Freising). NCBI’s RefSeq assembly GCF_000346065.1 was used for genome reference and annotation.

Illumina raw reads were preprocessed using Cutadapt (v3.5). Illumina TruSeq HT Index 1 (i7) adapters were removed from the 3’-end and nucleotides with a Phred quality score below 20 and their following downstream (5’–3’) bases were also removed. Then, the RNA-seq tool READemption (v2.0.4, DOI: https://doi.org/10.5281/zenodo.8421676 (Förstner et al. 2014)) was used to remove short reads (less than 20 nucleotides) and read mapping, gene quantification and differential gene expression analysis was conducted. Read mapping was performed using the aligner segemehl (v0.3.4) that is integrated into READemption, using a mapping accuracy of 95%. Differential gene expression was analyzed with DESeq2 (v1.34.0 (Love et al. 2014)) which is also part of READemption’s workflow. Differentially expressed genes (DEGs) were defined as log_2_(FC) > 1 and < -1 and p_adj_ ≤ 0.01. P-values were adjusted for multiple testing by DESeq2’s default Benjamini and Hochberg method. Differentially expressed genes were analyzed using gene set enrichment analysis and over-representation tests conducted with clusterProfiler (v. 4.10.1, R v. 4.3.2) (Wu et al. 2021), based on their associated GO and KEGG terms. The DEG detection was based on a set of *S. meliloti* RNA samples from ten conditions subjected to RNA-seq (see Data Availability statement) and used in the data analysis. DEG results were only reported for the comparison of *S. meliloti* 2011 cultures grown in TY medium and subjected to Tc exposure as specified above (Fig. 1A; Table S1).

For TSS detection, the RNA samples were split in two portions, and only one of them was treated with terminal exonuclease (TEX) (Sharma et al. 2010). The raw sequencing reads from TEX+/- libraries were preprocessed to remove Illumina TruSeq HT Index 1 (i7) adapters, and poly-A tails using Cutadapt (v3.5). As the sequencing was performed on a NextSeq 500 platform, which employs two-color chemistry, the nextseq=20 parameter was applied during quality trimming to account for platform-specific biases. Post-trimming, the quality of the reads was assessed using FastQC, ensuring data cleanliness and readiness for downstream analysis. READemption (v2.0.4) was used to map the trimmed reads. The resulting BAM files were processed to generate normalized coverage files. The normalized coverage was computed as the total number of aligned reads (abbreviated as tnoar) and then multiplied by the lowest number of aligned reads of all considered libraries. Subsequently, the normalized coverage files were used as input for prediction of TSS. This prediction was performed with default parameters using ANNOgesic (v1.1.14, https://doi.org/10.5281/zenodo.597066) (Yu et al. 2018), which employs TSSPredator as predictor. The entire bioinformatical workflow is published on Zenodo (https://doi.org/10.5281/zenodo.14671056). The workflow includes all necessary software in Docker containers, the entire input, results, custom Python and R scripts and an executable shell script that contains every command to ensure reproducibility.

For manual detection of TSSs upstream of DEGs, nucleotide-wise normalized coverage files from the TEX +/- libraries were visualized in the Integrated Genome Browser. Sharp coverage peaks representing 5’-ends of RNA, which were located up to 200 nt upstream of a start codon and were enriched or not diminished in the TEX-treated sample, were considered TSSs.

### Northern blot

For Northern blot hybridization, 10 µg RNA was separated in a 20×20 cm 10 % polyacrylamide-urea gel in TBE buffer at 300 V for 4 h. Transfer to a positively charged nylon membrane was conducted for 2 h at 150 mA using a Semi-Dry Blotter. The membrane was incubated at 56°C in pre-hybridisation solution (6× SSC, 0.5 % SDS, 10 µg/ml Salmon Sperm DNA, and 2.5× Denhardts solution) for 2 h and then overnight in hybridization solution (5 µl radioactively labelled oligonucleotide in 6× SSC, 0.5 % SDS and 10 µg/ml Salmon Sperm DNA). Thereafter, the membrane was washed in 0.01 % SDS, 5× SSC at room temperature. Signals were detected using a Bio-Rad molecular imager and Quantity One (Bio-Rad) software. The membrane was re-hybridized several times. The last probe was directed against 5S rRNA, which was used as loading control.

For radioactive labelling, 10 pmol of an oligonucleotide (Table S5) was mixed with 5′ phosphorylated by [^32^P]-adenosine-5′-triphosphate (3000 Ci/ mmol, 10 mCi/ml) and 5 U of T4-polynucleotid-kinase in buffer A (NEB). After incubation for 1 h at 37°C, 40 µl STE-buffer (10 mM Tris-HCl pH 8, 0.1 M NaCl, 2 M EDTA) was added to the 10 µl labelling reaction mixture to stop the reaction. G25-columns (GE Healthcare) were used to remove unbound nucleotides.

### qRT-PCR

Amount of mRNA was quantified using Brilliant III Ultra Fast SBR® Green QRT-PCR Mastermix (Agilent) and Luna Universal One-Step RT-qPCR Kit.

For determination of steady-state mRNA levels, first residual DNA was removed by treatment of 10 µg RNA with 1 µl TURBO-DNase (Invitrogen). The qRT-PCR was performed in a spectrofluorometric thermal cycler (Bio-Rad). BioRad CFX Manager 3.0 was used to determine the quantification cycle (Cq; the cycle, at which the amplification maximum of the curvature was reached) and the primer pair efficiencies in the respective PCR reaction. The Pfaffl-formula was used to calculate fold changes of mRNA amounts. The used reference mRNA was *rpoB*. Unless stated otherwise, the qRT-PCR analyses were conducted in three independent experiments with technical duplicates.

Determination of mRNA half-lives with qRT-PCR was performed as described (Scheuer et al. 2024). Briefly, culture samples were withdrawn 1, 2, 4, 6, and 8 min after rifampicin addition, RNA was isolated as described above together with a spike-in *rrp4* transcript, treated with TURBO-DNase and subjected to qRT-PCR analysis. The spike-in transcript was used as a reference. The RNA amount at the time point 0 was set to 100 % and linear-log graphs were used to calculate the mRNA half-lives. If the half-lives determined in three independent experiments were very different or no decay was detected in an experiment, a second primer pair targeting the same genes was used.

### 3’-RACE

DNA-free RNA isolated from non-treated, exponentially growing cultures, was depleted of rRNA using the NEBNext rRNA Depletion Kit for Bacteria (NEB). After rRNA depletion, the Universal miRNA Cloning Linker (NEB) was ligated to 3’ ends of transcripts using a T4 RNA ligase 2 (NEB). Then, RT-PCR was performed with primers (Table S5) specific to the adapter and the 5’ end of the *duf1127_1_* transcript. The amplicons were used as templates for a nested PCR. Since Taq polymerase was used, a blunting reaction was performed and the amplicons were cloned using the CloneJET PCR Cloning Kit (Thermo Scientific). Three biological experiments were performed and 24 cloned plasmids from each experiment were sequenced.

### Western blot analysis

Cells from 1.5 ml of a culture were harvested by centrifugation at 4°C and resuspended in 2x SDS buffer. Before loading on a 12% polyacrylamide gel for SDS-Tricine electrophoresis (Schägger 2006), samples were heated to 95°C for 10 min. Two gels were run in parallel using aliquots of the same samples. The one gel was used to transfer proteins were transferred to a polyvinylidene difluoride (PVDF) membrane with the help of a Semi-Dry Blotter, while the second gel was stained with Coomassie Brilliant Blue G-250 to control the protein amount of the samples. For detection of SPA-tag (Babu et al. 2012) containing proteins on the PVDF membrane, monoclonal, anti-FLAG tag antibodies conjugated with horseradish peroxidase (Sigma Aldrich) and Western Lightning Ultra, Chemiluminescent Substrate (PerkinElmer) were used. Signals were visualized with a Fusion-SL chemiluminescent imager (Peqlab).

### Fluorescence measurement

Aliquots (150 µl) of cultures of *S. meliloti* strains containing *egfp* reporter plasmids were transferred to a 96-well microtiter plate. Fluorescence (extinction at 488 nm and emission at 522 nm) and ODs of the samples were measured in a Tecan Infinite M200 reader. Fluorescence values were normalized to the measured OD_600 nm_ and the autofluorescence of an empty vector control culture. Three independent cultures technical triplicates were used for the measurements.

### Statistical analysis and RNA structure prediction

If not stated otherwise, all data are shown in means and single data points of three independent experiments. Technical duplicates were used in qRT-PCR and technical triplicates in fluorescence measurements. Statistical analysis including t-tests were performed in R (https://www.r-project.org/). For RNA structure prediction, RNAfold was used (Gruber et al. 2008).

## Author contributions

Conceptualization, E.E.H., J.K., K.U.F.; Methodology, J.K., T.S.; Investigation, J.K., T.S., R.S., T.D., J.W., S.B.W.; Formal analysis, J.K., T.S., R.S., T.D.; Writing – Original Draft, E.E.H., J.K.; Writing – Review and Editing, E.E.H., J.K., T.D., K.U.F.; Visualization, E.E.H., J.K., R.S., T.D.; Supervision, E.E.H., K.U.F.; Funding Acquisition, E.E.H.

## Acknowledgements

This work was funded by the Deutsche Forschungsgemeinschaft (Ev42-6/2; GRK2355, project number 62202628).

## Data Availability statement

The raw read files presented in this article have been submitted to the European Nucleotide Archives (https://www.ebi.ac.uk/ena/browser/view/PRJEB83671) under the accession number PRJEB83671. The authors confirm that the data supporting the findings of this study are available within the article and its supplementary materials.

