## Supplemental Figures 1 to 8 for "Early posttranscriptional response to tetracycline exposure in a gram-negative soil bacterium reveals unexpected attenuation mechanism of a DUF1127 gene"

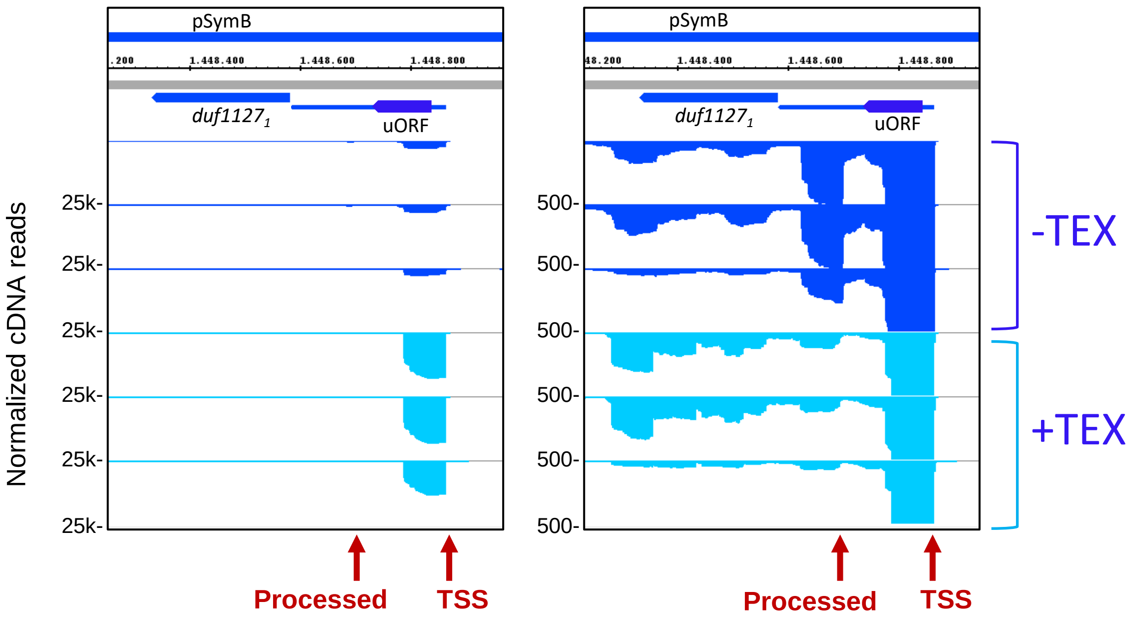


**Figure S1 Integrated genome browser view of the DUF1127$_1$ operon showing the transcriptional start site (TSS) upstream of the uORF and the processing site in the intergenic region (IGR) between the uORF and duf1127$_1$.** Shown are normalized cDNA reads of three biological experiments. The RNA sample from each experiment was split in two halves. The one of them was treated with a terminal exoribonuclease, which degrades processed transcripts lacking a triphosphate at their 5'-ends (+TEX), while the other one was not processed with TEX (-TEX). In the (+TEX) samples, the peak corresponding to the processed 5'-end in the IGR (marked with a vertical arrow labelled with proc.) was diminished (see the right panel), while the peak corresponding to the TSS was enriched (see the left panel).

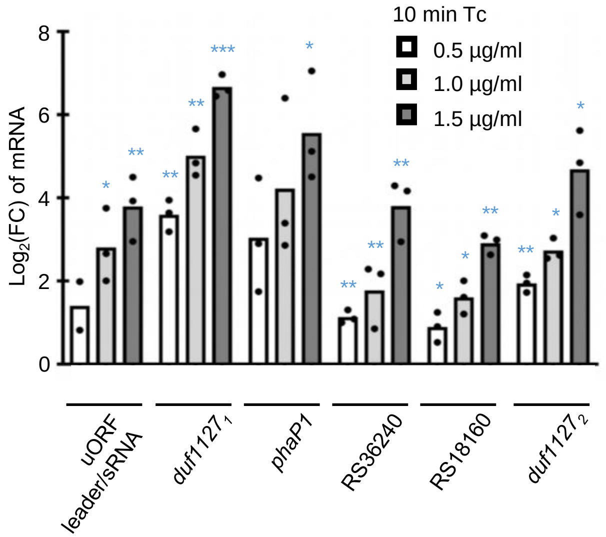


**Figure S2 Low Tc amounts lead to significant mRNA increase in 10 min**. Analysis by qRT-PCR of RNA level level changes 10 min after addition of tetracycline (Tc) at the indicated final concentrations. The analyzed mRNAs are indicated. All graphs show means and single data points of three independent experiments. *** $p \leq 0.001$, ** $p \leq 0.01$, * $p \leq 0.05$.

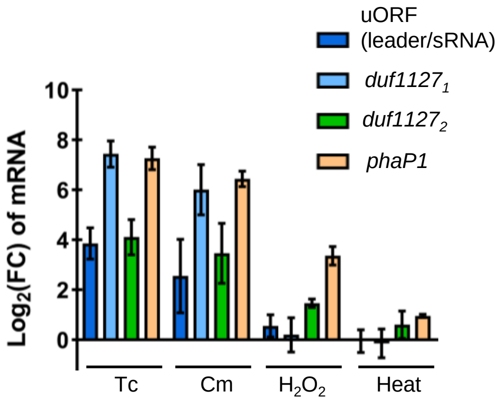


**Figure S3 The *duf1127$_1$* operon shows mRNA increase upon translation inhibition, but not upon oxidative and heat stress**. Analysis by qRT-PCR of RNA level changes upon 10 min of exposure to the indicated stressors. Tc: tetracycline, Cm: chloramphenicol. The analyzed mRNAs are indicated. All graphs show means and standard deviations of three independent experiments.

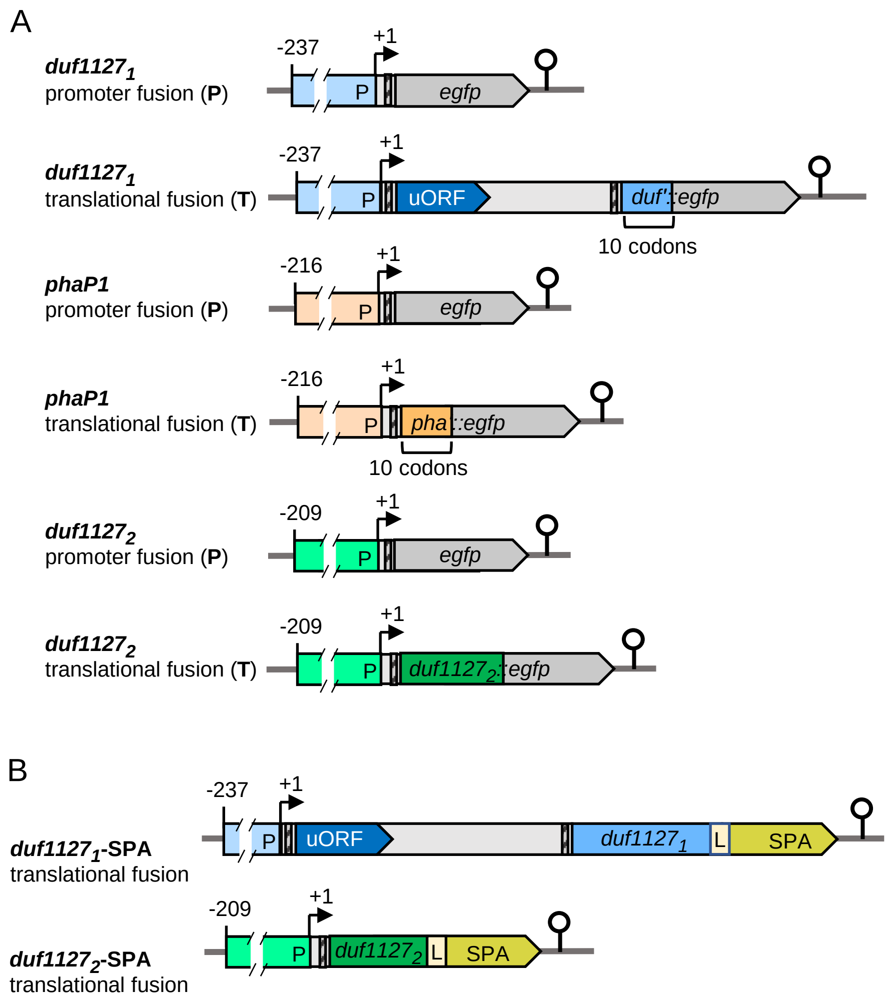


**Figure S4**

**Scheme of reporter fusions with *egfp* (A) and SPA-fusions of full-length *duf1127*$_1$ and *duf1127*$_2$ (B).**

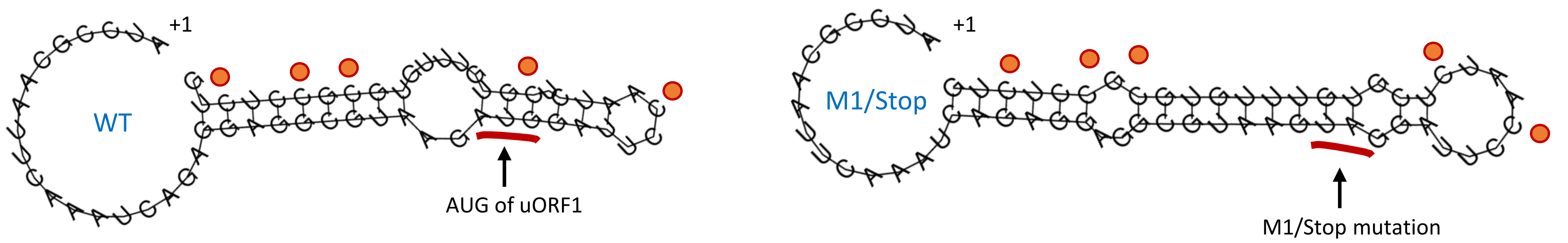


**Figure S5 RNA structure prediction of the 5'-region of the uORF1 transcript for the wild (WT) type sequence and the sequence harboring the M1/Stop mutation.** ● Cytidines subjected to exchange in the syn. mut. derivative of pDUF'-SPA (see Fig. 7).
